# Efficient toolkit implementing best practices for principal component analysis of population genetic data

**DOI:** 10.1101/841452

**Authors:** Florian Privé, Keurcien Luu, Michael G.B. Blum, John J. McGrath, Bjarni J. Vilhjálmsson

## Abstract

Principal Component Analysis (PCA) of genetic data is routinely used to infer ancestry and control for population structure in various genetic analyses. However, conducting PCA analyses can be complicated and has several potential pitfalls. These pitfalls include (1) capturing Linkage Disequilibrium (LD) structure instead of population structure, (2) projected PCs that suffer from shrinkage bias, (3) detecting sample outliers, and (4) uneven population sizes. In this work, we explore these potential issues when using PCA, and present efficient solutions to these. Following applications to the UK Biobank and the 1000 Genomes project datasets, we make recommendations for best practices and provide efficient and user-friendly implementations of the proposed solutions in R packages bigsnpr and bigutilsr.

For example, we find that PC19 to PC40 in the UK Biobank capture complex LD structure rather than population structure. Using our automatic algorithm for removing long-range LD regions, we recover 16 PCs that capture population structure only. Therefore, we recommend using only 16-18 PCs from the UK Biobank to account for population structure confounding. We also show how to use PCA to restrict analyses to individuals of homogeneous ancestry. Finally, when projecting individual genotypes onto the PCA computed from the 1000 Genomes project data, we find a shrinkage bias that becomes large for PC5 and beyond. We then demonstrate how to obtain unbiased projections efficiently using bigsnpr.

Overall, we believe this work would be of interest for anyone using PCA in their analyses of genetic data, as well as for other omics data.

## 1 Introduction

Principal Component Analysis (PCA) has been widely used in genetics for many years and in many contexts. For instance, adding PCs as covariates is routinely used to adjust for population structure in Genome-Wide Association Studies (GWAS) (Price *et al.* 2006; Novembre and Stephens 2008). PCA has also been used to detect loci under selection (Galinsky *et al.* 2016; Luu *et al.* 2017) and in heritability analyses Yang *et al.* (2010); Loh *et al.* (2015a). Recently, the advent of large population-scale genetic datasets, such as the UK biobank data, has prompted research on developing scalable algorithms to compute PCA on very large data (Bycroft *et al.* 2018). It is now possible to efficiently approximate PCA on very large datasets thanks to software such as FastPCA (fast mode of EIGENSOFT), FlashPCA2, PLINK 2.0 (approx mode), bigstatsr/bigsnpr, TeraPCA and ProPCA (Galinsky *et al.* 2016; Abraham *et al.* 2017; Chang *et al.* 2015; Privé *et al.* 2018; Bose *et al.* 2019; Agrawal *et al.* 2019).

However, in practice, conducting PCA on genotype data to capture population structure consists of more steps than simply performing singular value decomposition on the genotype matrix. These includes removing related individuals, pruning variants in LD, and excluding outlier samples that can suggest poor genotyping quality. Indeed, many pitfalls related to PCA of genotype data have been documented and none of the currently available software address all of these. In the following, we outline these pitfalls and explain when they are relevant. First, some of the PCs may capture LD structure rather than population structure (Price *et al.* 2008; Abdellaoui *et al.* 2013; Privé *et al.* 2018). Including PCs that capture LD as covariates in genetic analyses can lead to reduced power for detecting genetic association within these LD regions. Second, another issue may arise when projecting PCs of a reference dataset to another study dataset: projected PCs are shrunk towards 0 in the new dataset (Lee *et al.* 2010; Wang *et al.* 2015; Zhang *et al.* 2019). This shrinkage makes it potentially dangerous to use the projected PCs for analyses such as PC regression, ancestry detection and correction for ancestry. This is also an issue when projecting individuals from the same dataset that were discarded from the PCA computation (e.g. related individuals). Third, PC scores may capture outliers that are due to family structure, population structure or other reasons; it might be beneficial to detect and remove these individuals to maximise the population structure captured by PCA (in the case of removing a few outliers) or to restrict analyses to genetically homogeneous samples (e.g. “White British” people in the UK Biobank). Finally, efficient methods for PCA use approximations, which can results in some lack of precision of computed PCs. This has been shown to be a possible issue for software such as FastPCA and PLINK 2.0, but not for FlashPCA2 and bigstatsr/bigsnpr (Abraham *et al.* 2017; Privé *et al.* 2018). An overview of existing methods with their respective advantages and limitations is presented in table 1.

**Table 1:**
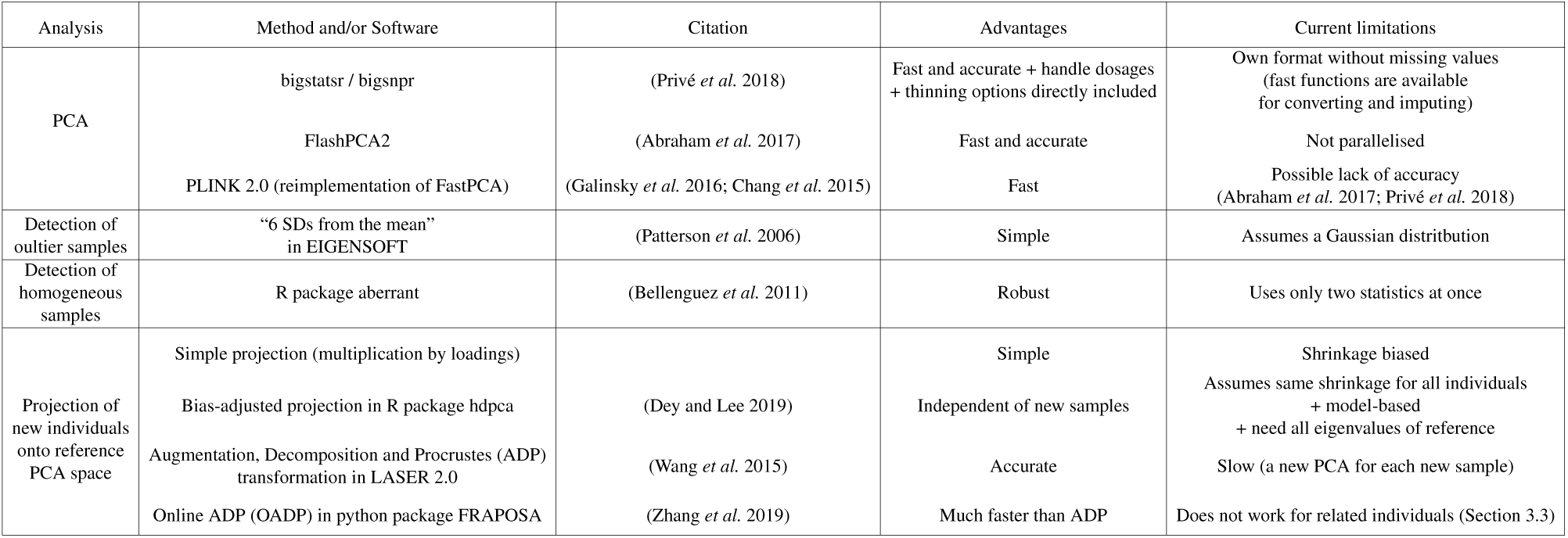
Overview of existing methods.

For this paper, we derive implementations of truncated PCA and other useful functions for e.g. performing LD thinning and computing various statistics. We make these available in a new release of R package bigsnpr (v1.0.0); what differs from previously available functions presented in Privé *et al.* (2018) is that these new functions can be used directly on PLINK bed/bim/fam files with some missing values. We use these new functions to analyse the UK Biobank data, and show that they are both very fast and easy to use. We also point out that many PCs currently reported by the UK Biobank capture LD structure instead of population structure. Interestingly, subsetting the UK Biobank data enables to get more PCs that capture population structure than when using the whole sample (∼40 instead of ∼16). Then, we project the other individuals that were not used in the PCA calculation, show that this projection is biased and provide an efficient solution to estimate unbiased projections instead. Finally, we explore options to detect outlier samples in PCA, either a few outlier samples that may correspond to e.g. batch effects or family structure, or when the goal is to restrict the data to individuals of homogeneous ancestry.

## 2 Material and Methods

### 2.1 Efficient implementation of PCA for genotype data

When there is no missing value, we compute the partial Singular Value Decomposition (SVD) *U* Δ*V* ^*T*^ of the scaled genotype matrix of diploid individuals 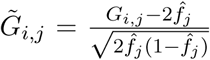 where *G*_*i,j*_ is the allele count (genotype) of individual *i* and variant *j*, and 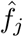 is the estimated allele frequency of variant *j* (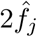 is the mean allele count of variant *j*). Then, *U* Δ are the first *K* PC scores and *V* are the first *K* PC loadings, where *K* is the number of PCs computed (e.g. *K* = 20).

When there are some missing values, we compute the partial SVD similarly, except that missing values are replaced by the variant means (i.e. 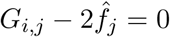 when *G*_*i,j*_ is missing) and the 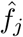 for each variant are estimated using only non-missing values of each variant. Note that this decomposition is equivalent to the decomposition presented above after imputation by the variant means.

To compute this decomposition easily and efficiently, we implement an accessor that memory-map the PLINK bed file to use it directly as if it were a standard matrix. Then, we apply the same algorithm for partial SVD that is used in R packages bigstatsr and FlashPCA2, namely the implicitly restarted Arnoldi method (Lehoucq and Sorensen 1996; Abraham *et al.* 2017; Privé *et al.* 2018). This algorithm is implemented in R package RSpectra, and requires a function that computes the matrix-vector multiplication of the scaled genotype matrix with a given vector, which we implement for PLINK bed files and parallelise.

### 2.2 Robust Mahalanobis distance

Mahalanobis distances are computed as *d*(*x*)^2^ = (*x*−*µ*)^*T*^ Σ^−1^(*x*−*µ*), where *µ* and Σ are (robust) estimators of location and covariance of *x*. We use these distances for many applications in this paper. When *x* is multivariate Gaussian data with *K* dimensions, the squared distances follow a *χ*^2^(*K*) distribution. If *x* are PC scores of centered data and if we use standard estimates, then *µ* = 0 and Σ = *I*_*K*_. Yet, here we use the pairwise orthogonalised Gnanadesikan-Kettenrin robust estimates of these parameters, as implemented in function `covRob` of R package robust with parameter `estim = “pairwiseGK”` (Gnanadesikan and Kettenring 1972; Yohai and Zamar 1988; Maronna and Zamar 2002; Todorov *et al.* 2009). We reexport function `covRob` in R package bigutilsr for convenience.

### 2.3 Detecting LD structure in PCA

For detecting outlier variants in PCA that are due to long-range Linkage Disequilibrium (LD) regions, we use a similar procedure as described by Privé *et al.* (2018). We first apply a first round of clumping at e.g. *r*^2^ > 0.2, prioritising variants by higher minor allele count. Then, we compute *K* PC scores and loadings. To summarise the contribution of each variant in all *K* PC loadings, we compute the robust Mahalanobis distances (Section 2.2) of these PC loadings. To capture consecutive outliers that corresponds to long-range LD regions, we apply a Gaussian smoothing to these statistics (moving average with a Gaussian filter over a window of 50 variants as radius by default).

Finally, to choose the threshold on the previously described statistics above which variants are considered outliers, we use a modified version of Tukey’s rule, a standard rule for detecting outliers (Tukey 1977). The standard upper limit defined by Tukey’s rule is *q*_75%_(*x*) + 1.5 · *IQR*(*x*), where *x* is the vector of computed statistics and *IQR*(*x*) = *q*_75%_(*x*)−*q*_25%_(*x*) is the interquartile range. One assumption of Tukey’s rule is that the sample is normally distributed; we account for skewness in the data using the medcouple as implemented in function `adjboxStats` of R package robustbase (Brys *et al.* 2004; Hubert and Vandervieren 2008). Standard Tukey’s rule also uses a fixed coefficient (1.5) that does not account for multiple testing, which means that there are always some outliers detected when using 1.5 for large samples. To solve these two potential issues, we implement `tukey_mc_up` in R package bigutilsr and use it here, which accounts both for skewness and multiple testing by default.

We remove the detected outlier variants, compute the PC scores and loadings again, and iterate until there is no detected outlier variant anymore. This procedure is implemented in function `bed_autoSVD` of R package bigsnpr.

### 2.4 Detecting outlier samples in PCA

For detecting outlier samples in PCA, we use a modified version of the Probabilistic Local Outlier Factor (PLOF) statistic on PCs (Kriegel *et al.* 2009). Using K nearest neighbours (KNN), this consists in comparing the distance from a point *j* to its KNNs (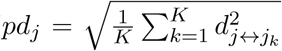, where *j*_k_ is the *k*-th NN of *j*) with the distances from its KNNs to their respective KNNs 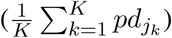. The idea is that an outlier should be far from all other points, and is even more outlier if its KNNs are in a very dense cluster. Here, we use 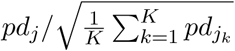. Note the square root, as it otherwise detects as outlier any point that is next to a very dense cluster. We implement (the two parts of) this statistic in function `prob_dist` of R package bigutilsr. To make it fast, we use the fast K nearest neighbours implementation of R package nabor (Elseberg *et al.* 2012) and parallelise it in function `knn_parallel` of package bigutilsr. Automatic outlier detection is difficult, therefore we recommend users to choose a threshold for this statistic to define outliers based on visual inspection (using the histogram of these statistics and the PC scores colored by these statistics; see e.g. figure 2).

As for detecting samples that have a different ancestry from most of the samples in the data, i.e. for restricting to homogeneous samples, we compute the pairwise orthogonalised Gnanadesikan-Kettenrin robust Mahalanobis distances on PC scores (Section 2.2). We then restrict to individuals whose log-distance (alternatively p-value) is smaller (larger) than some threshold determined based on visual inspection.

### 2.5 Projecting PCs from a reference dataset

To project PCs of a reference dataset (e.g. 1000G) to a target genotype dataset, we implement the following 3 steps in function `bed_projectPCA` of package bigsnpr: 1) matching the variants of each dataset, including removing ambiguous alleles [A/T] and [C/G], and matching strand and direction of the alleles; 2) computing PCA of the reference dataset using the matched variants only; 3) projecting computed PCs to the target data using an optimised implementation (see Supplementary Materials) of the Online Augmentation, Decomposition, and Procrustes (OADP) transformation (Zhang *et al.* 2019). To project individuals from the same dataset as the ones used for computing PCA, you can use function `bed_projectSelfPCA`. Note that the new individuals to be projected should not be related to the ones used for computing PCA (Section 3.3).

### 2.6 Data

We provide and use a subsetted version of the 1000 genomes (1000G) project data (1000 Genomes Project Consortium *et al.* 2015; Meyer 2019). Variants are restricted to the ones in common with HapMap3 or UK Biobank (International HapMap 3 Consortium *et al.* 2010; Bycroft *et al.* 2018). Moreover, we apply some quality control filters; we remove variants having a minor allele frequency < 0.01, variants with P-value of the Hardy-Weinberg exact test < 10^−50^, and non-autosomal variants. To remove related individuals with second-degree relationship or more, we apply KING-relatedness cutoff of 0.0884 to the data using PLINK 2.0 (Manichaikul *et al.* 2010; Chang *et al.* 2015). This results in 2490 individuals and 1,664,852 variants of the 1000G project (phase 3) in PLINK bed/bim/fam format. Resulting PLINK files and R code to generate these files are made available at https://doi.org/10.6084/m9.figshare.9208979.v3. To easily download these data, we provide function `download_1000G` in R package bigsnpr.

In this paper, we also analyse the UK Biobank data (https://www.ukbiobank.ac.uk/). We apply some quality control filters; we remove individuals with more than 10% missing values, variants with more than 1% missing values, variants having a minor allele frequency < 0.01, variants with P-value of the Hardy-Weinberg exact test < 10^−50^, and non-autosomal variants. This results in 488,371 individuals and 504,139 variants. When removing related individuals, we use the list of individual pairs reported by the UK Biobank.

## 3 Results

### 3.1 Application to the UK Biobank

To demonstrate that we provide very fast implementations of the different methods presented in this paper, we apply them to the UK Biobank. We use 20 physical cores for most of the computations (CPU: Intel(R) Xeon(R) Silver 4114, 2.20GHz). It takes 22 minutes to perform a first phase of clumping on 406,545 unrelated individuals genotyped over 504,139 variants, which reduces the number of variants to 261,307. It then takes 34 minutes to compute the first 20 PCs using these 261,307 variants. When performing the automatic procedure for LD detection, it takes 5 hours to perform the initial clumping step, 6 rounds of computation of PCs and 5 rounds of outlier variant detection (i.e. 5 iterations of outlier detection and one final computation of PCs).

When applying our automatic procedure to remove long-range LD regions, it does not converge after 5 iterations for the UK Biobank, meaning that it keeps detecting long-range LD regions at each iteration (represented by peaks in PC loadings). Therefore, we are able to capture only 16 PCs that show stratification that is not LD structure (Figures S8, S9 and S10). Similarly, PC loadings reported by the UK Biobank clearly show that PC19 to PC40 capture LD structure, which is also the case for PC16 and PC18, although less pronounced (see peaks in figure 1). These include e.g. one region on chromosome 6 (70-91 Mbp) that is captured in PC19 (Figure S10) and that was not previously reported in Price *et al.* (2008).

**Figure 1:**
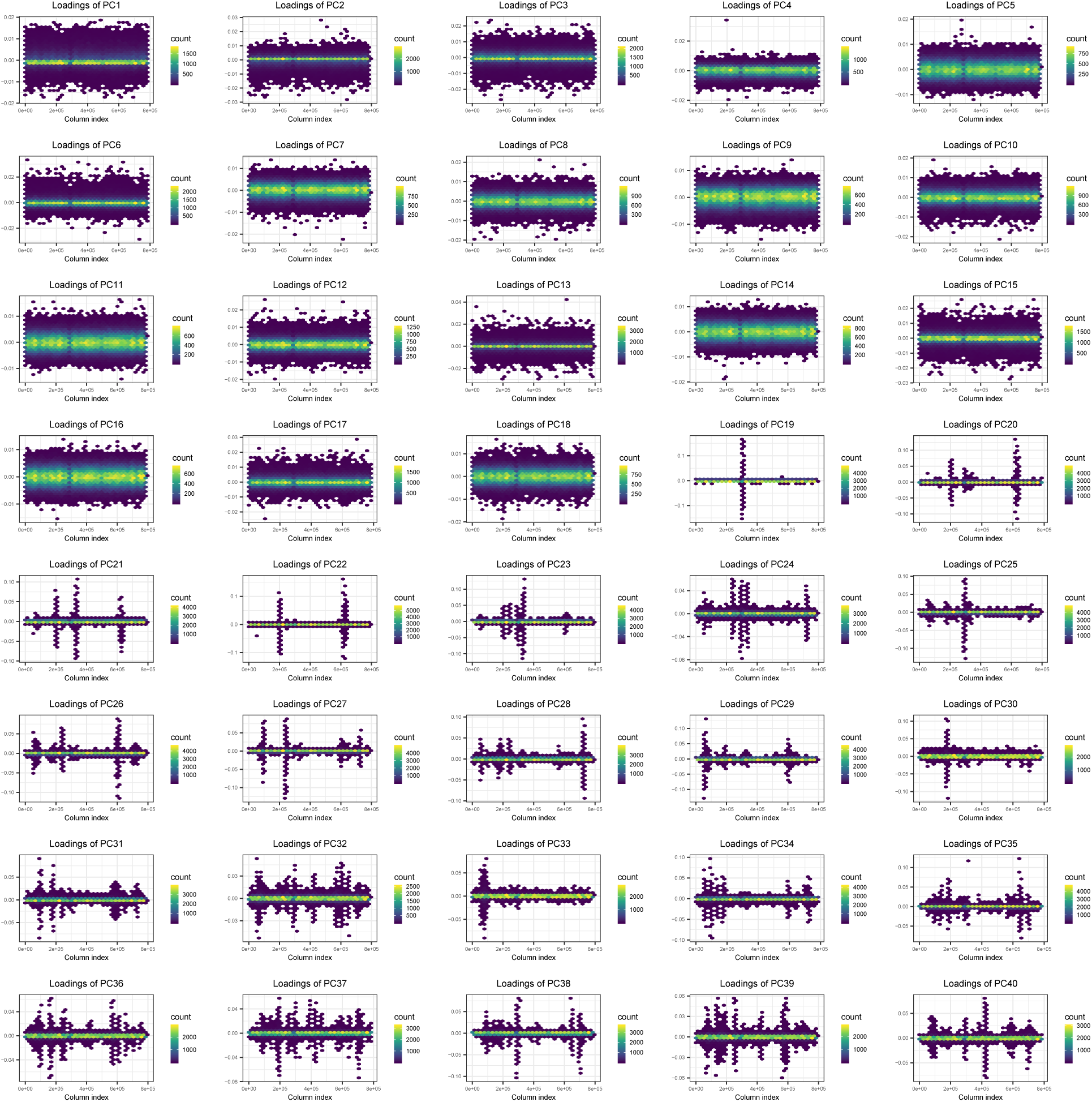
Principal Component (PC) loadings 1 to 40 reported by the UK Biobank. X-axis represent positions of variants in the data, which is ordered by chromosome and physical position, and y-axis the value of loadings. Points are hex-binned to make plotting of such large number of points manageable.

As for other analyses, it takes 8 minutes to match the 1000G data to the UKBB data and compute 20 PCs of the 1000G data using the automatic LD detection technique. It takes 12 minutes more to do the OADP projection of all 488,371 individuals of the UKBB data onto the PCA computed using the 1000G data. Finally, it takes only 6 minutes to compute the 30-Nearest Neighbours of 20 PC scores for 406,545 UK Biobank individuals, which is the main computational aspect of computing the statistics used to detect individual outlier samples (Section 2.4).

### 3.2 Outlier sample detection

To detect a few sample outliers, we compare the standard rule of “6 SDs from the mean” (6SD) used in e.g. EIGENSOFT to the statistic we propose in section 2.4. Our statistic identifies only isolated samples or isolated pairs that seems to be outliers driving structure of PC17-20 of 1000G (Figure 2). All but one outliers are distantly-related pairs that disappear if using a more stringent threshold on relatedness (i.e. using a KING-relatedness cutoff of ∼0.0442 instead of ∼0.0884, see tutorial in section “code availability”). In contrast, rule 6SD identifies a lot of outliers, of which some are part of a relatively large cluster (Figure S1). We recall that, in theory, if all PCs are normally distributed, after correcting for multiple testing of 2500 individuals and 20 PCs, 6SD has a small probability of 0.0001 of detecting one outlier or more.

**Figure 2:**
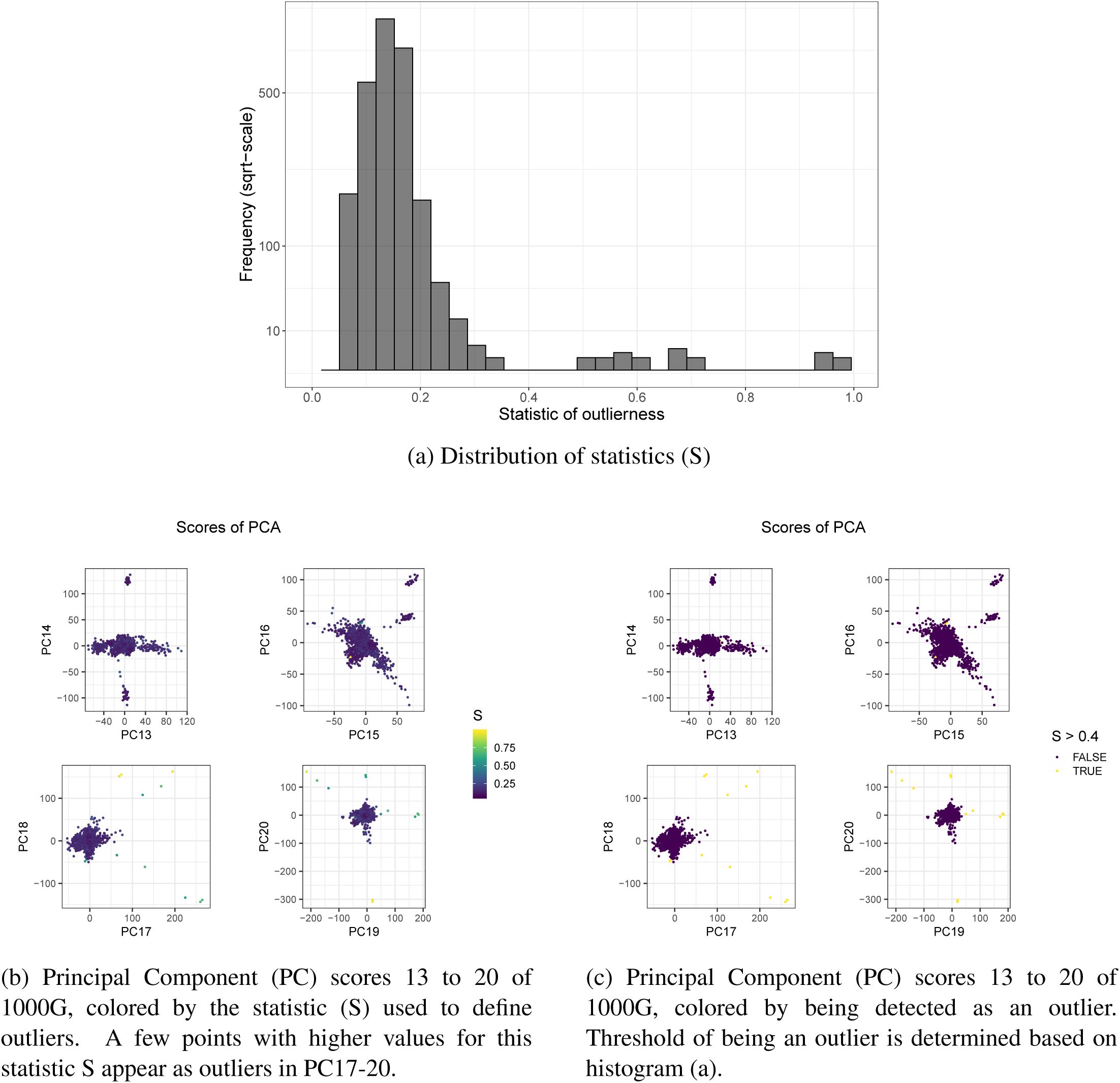
Outlier detection in the 1000 Genomes (1000G) project, using `prob_dist` (Section 2.4).

As for restricting to homogeneous samples, we compare the use of R package aberrant used to report the “White British” subset of UKBB homogeneous samples, to the use of the robust Mahalanobis distance we propose. Here, we visually choose a threshold of 5 on the log-distance and show that this gives a similar subset of individuals than the “White British” subset reported by the UK Biobank (Figure S2). Moreover, when using this threshold, only 3 out of 10,936 people of self-reported Asian ancestry (1 “Chinese” and 2 “Indian”) are kept, and 1 “African” out of 7622 people with Black background is kept (Table S1). In contrast, 416,492 out of 431,090 “British” (96.6%) and 12,620 out of 12,759 “Irish” (98.9%) are kept.

### 3.3 Projecting onto the PCA space from a reference dataset

We use 60% of individuals in the 1000G data (section 2.6) to compute K=20 PCs. Then, we project the remaining 40% individuals using three methods: 1/ simply multiplying the genotypes of these individuals by the previously computed loadings; 2/ correcting the simple projections using asymptotic shrinkage factors as determined by R package hdpca v1.1.3 (Dey and Lee 2019), with all eigenvalues derived from the Genetic Relationship Matrix (GRM) computed with `bed_tcrossprodSelf`, one of the new functions of R package bigsnpr; and 3/ the OADP projection (section 2.5). When simply projecting using loadings, there is negligible shrinkage for PC1 and PC2, a small shrinkage for PC3 and PC4, and a large shrinkage for PC5 to PC8 (Figure 3). In contrast, there is no visible shrinkage when projecting new individuals with OADP (Figure 3). Simple projection is affected even more by this shrinkage for PC9 to PC20, while OADP still appears free of this bias (Figure S3). We show the same results when projecting the full UK Biobank data onto PCA computed using 1000G data (Figure S6). When correcting projected PC scores with asymptotic shrinkage factors, bias is smaller than with simple projection, yet, there is a visible bias for PC7-8 (Figure S4). Finally, to assess if OADP could be used to project individuals that are related to some individuals that were used to compute PCA, we projected these 60% individuals (as if we were projecting their monozygotic twins) using OADP. Projections of related individuals using OADP shows some bias in reverse direction (Figure S5).

**Figure 3:**
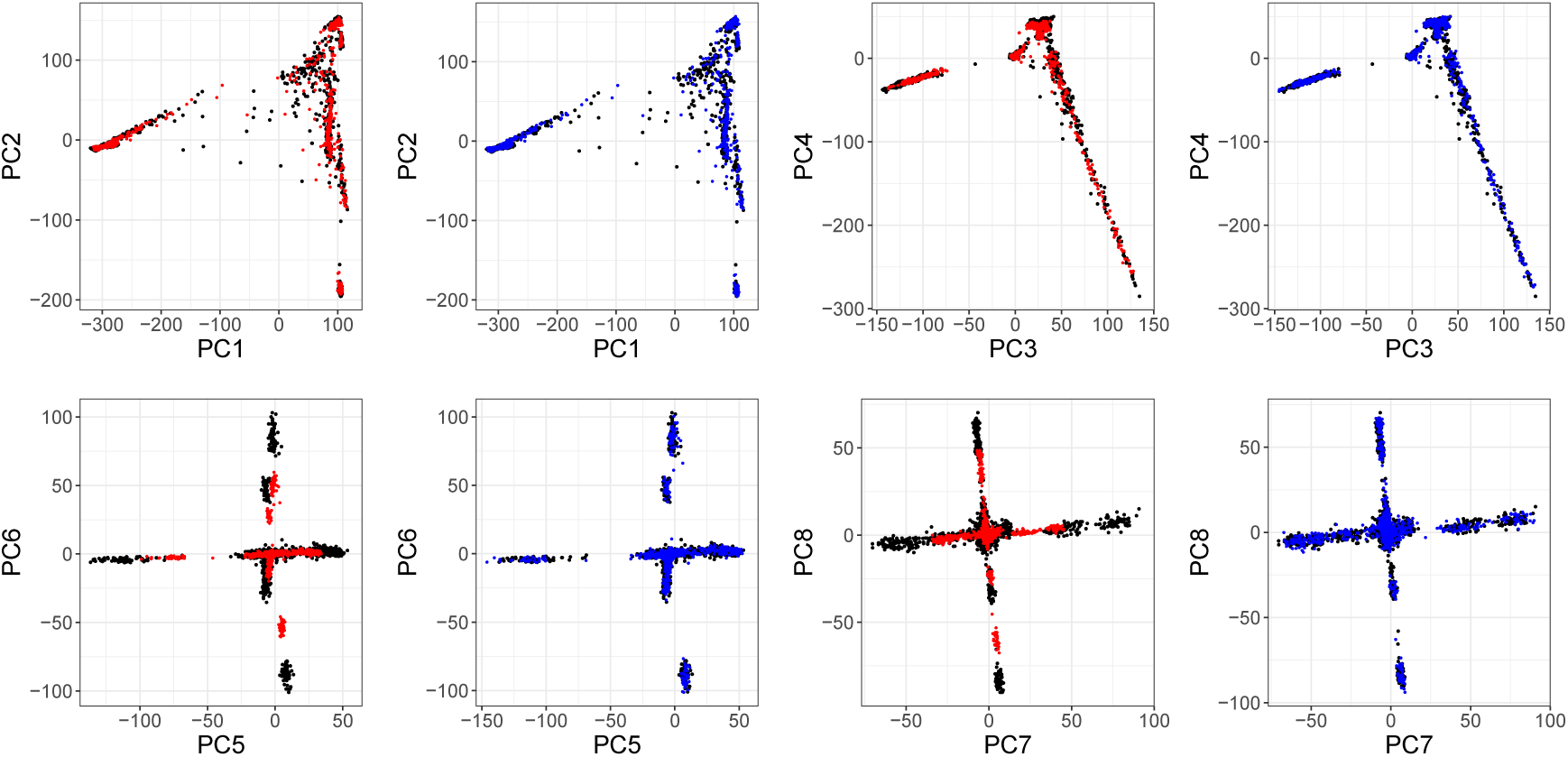
Principal Component (PC) scores 1 to 8 of the 1000 Genomes project. Black points are the 60% individuals used for computing PCA. Red points are the 40% remaining individuals, projected by simply multiplying their genotypes by the corresponding PC loadings. Blue points are the 40% remaining individuals, projected using the Online Augmentation, Decomposition, and Procrustes (OADP) transformation. Estimated shrinkage coefficients for these 8 PCs are 1.01 (PC1), 1.02, 1.06, 1.09, 1.50 (PC5), 1.69, 1.98 and 1.39.

When computing the PCs on the UK Biobank using 406,545 unrelated individuals and 171,977 variants, and projecting the 1000G data onto this reference PCA space, shrinkage is much smaller (≤ 1.08 for all 20 first PCs, Figure S7). Overall, this shrinkage decreases with an increased sample size (Table 2).

**Table 2:** Shrinkage coefficients when projecting new individuals onto reference PCA space. We list the dataset, the sample size and number of variants used to compute the final PCA. As expected, the shrinkage bias only becomes negligible if the PCA is conducted on large samples.

| Dataset | Sample size ($\times 1000$) | Number of variants<br>($\times 1000$, after LD removal) | Shrinkage (PC 1 - 5 - 10 - 20) |
| --- | --- | --- | --- |
| 1000G | 1.5 | 393 | 1.01 - 1.50 - 3.14 - 6.70 |
| 1000G | 2.5 | 229 | 1.01 - 1.36 - 2.84 - 6.75 |
| UKBB | 49.0 | 282 | 1.00 - 1.04 - 1.12 - 1.43 |
| UKBB | 406.5 | 172 | 1.00 - 1.01 - 1.04 - 1.08 |

### 3.4 Capturing subtle population structure in the UK Biobank

We recomputed PCA in the UK Biobank after restricting the individuals included in the computations: we randomly subsampled UKBB data to use only 10,000 British individuals (out of 431,029) and 5000 Irish individuals (out of 12,755), while keeping all individuals with other or unknown self-reported ancestry. We further removed all pairs of related individuals reported by the UKBB (i.e. both individuals in each pair). This resulted in 48,942 individuals that we used to compute 50 PCs, which took less than 3 hours using function `bed_autoSVD` (that converged after 4 iterations of automatic LD removal). We show that we are able to capture more PCs (at least 40 instead of 16-18) that display visual population structure (Figures 4 and S12). We then projected all 439,429 remaining individuals from UKBB onto this PCA space in 21 minutes only using our implementation of the OADP projection (`bed_projectSelfPCA`). Note that these individuals should not be related to any of the 48,942 individuals used for training PCA because we removed both individuals from each pair of related individuals in UKBB. Projection of new individuals show again a clear shrinkage when using simple projection (between 1.00 for PC1 and 1.80 for PC50), but no visible bias when using OADP projection (Figure S13).

**Figure 4:**
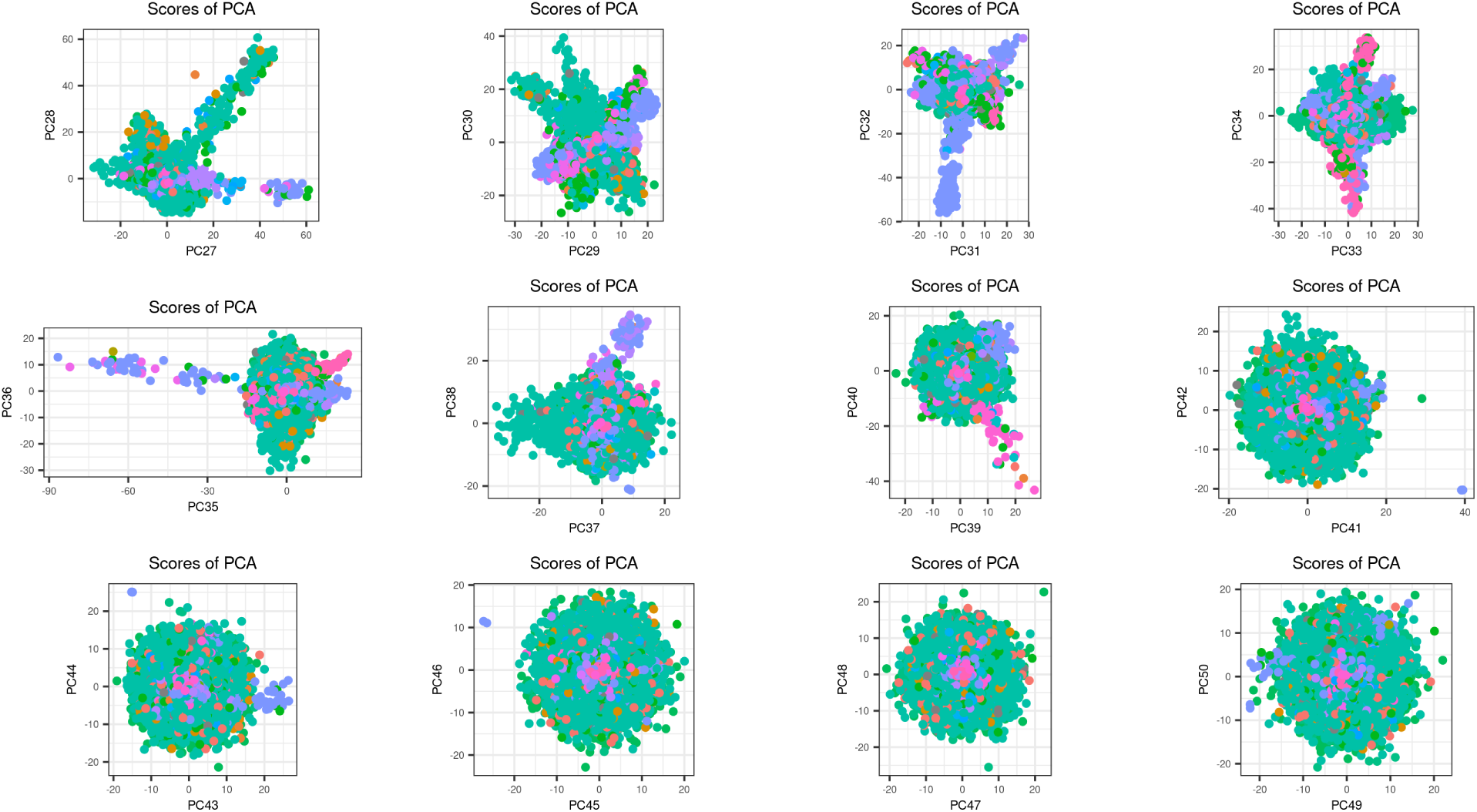
Principal Component (PC) scores 27 to 50 computed on the UK Biobank using 48,942 individuals of diverse ancestries. These individuals are the ones resulting from removing all related individuals and randomly subsampling the British and Irish individuals. Different colors represent different self-reported ancestries.

### 3.5 PCA & missing value imputation

As we compute PCA on data with missing values, although we restrict to variants with less than 1% missing values, we analyse hereinafter the effect of imputation of missing values before computing PCA. We compare 4 different imputation methods and two different sets of individuals. In the UK Biobank imputed data, approximately 1000 individuals have been removed because of a high number of missing values or a high heterozygosity, as compared to the genotyped data (Bycroft *et al.* 2018). When computing PCA with mean imputation and using all genotyped individuals, PC16 is capturing individuals with very high heterozygosity (Figure S11). When restricting to imputed individuals only, i.e. after removing individuals with very high heterozygosity, PC16 completely disappears and new PC16 to PC19 correspond to previous PC17 to PC20 (Table S2). When using dosage data instead of genotype data with mean imputation, PCA is globally unchanged (Table S3). Overall, if we choose to use either one of the following 4 imputation methods: mean imputation, random imputation according to allele frequencies, using reported dosage data from BGEN files, or imputation of genotyped data based on machine learning using function `snp_fastImpute` of R package bigsnpr (Privé *et al.* 2018), resulting PCs are always very similar (absolute correlation larger than 0.99 for the 20 computed PCs, results not shown). This justifies performing PCA with mean imputation directly on PLINK bed files with a few missing values; this has the advantage to be much faster than having to impute genotyped data using `snp_fastImpute`, which took 4 days for 406,545 individuals and 240,444 variants, or based on external reference datasets.

## 4 Discussion

In this work, we have compiled different pitfalls that can arise with Principal Component Analysis (PCA) of genetic data. Then, we have investigated possible solutions to these pitfalls and selected the ones that we found most advantageous, both with respect to properties such as accuracy and robustness, but also computational efficiency and ease of use. We then implemented these solutions in R packages bigsnpr and bigutilsr. The new functions we provide in R package bigsnpr can be directly applied to genotypes stored as PLINK bed/bim/fam files with some missing values. This contrasts with previous releases of bigsnpr that could only use format “bigSNP”. This data format can store both genotype calls and dosages, but requires conversion from other formats and imputation of missing values using functions provided in the package (Privé *et al.* 2018). As PCA is a useful tool on its own and does not require extensive imputed data, we therefore decided that operating directly on PLINK files with a few missing values would be more practical for users.

We summarise our work into several recommendations for computing PCA, and propose the pipeline shown in figure 5. Note that we have not included standard steps such as initial quality control filters and post-analysis checks (e.g. visual inspection of different plots). This pipeline requires removing all related individuals, for which we provide an R wrapper to PLINK’s implementation of KING robust kinship coefficients (Manichaikul *et al.* 2010; Chang *et al.* 2015). Note that one should remove both individuals in each pair of related individuals. This ensures that the projected individuals are not related to the ones used for computing PCA, since we showed that relatedness is a problem when using the OADP projection (Figure S5). After selecting a subset of individuals, we apply several steps of outlier detection, one for outlier variants that capture long-range LD variation (automatic), and one for detecting outlier samples (semi-automatic and visual). To make these steps more computationally efficient, we explored solutions for not recomputing PCA from scratch when removing a few samples or a few variants. Using educated guesses in R package PRIMME based on low-rank approximations of the updated PCA seemed to be a promising approach but did not reduce computation time by much, so we did not pursue this idea (Brand 2003; Wu *et al.* 2017).

**Figure 5:**
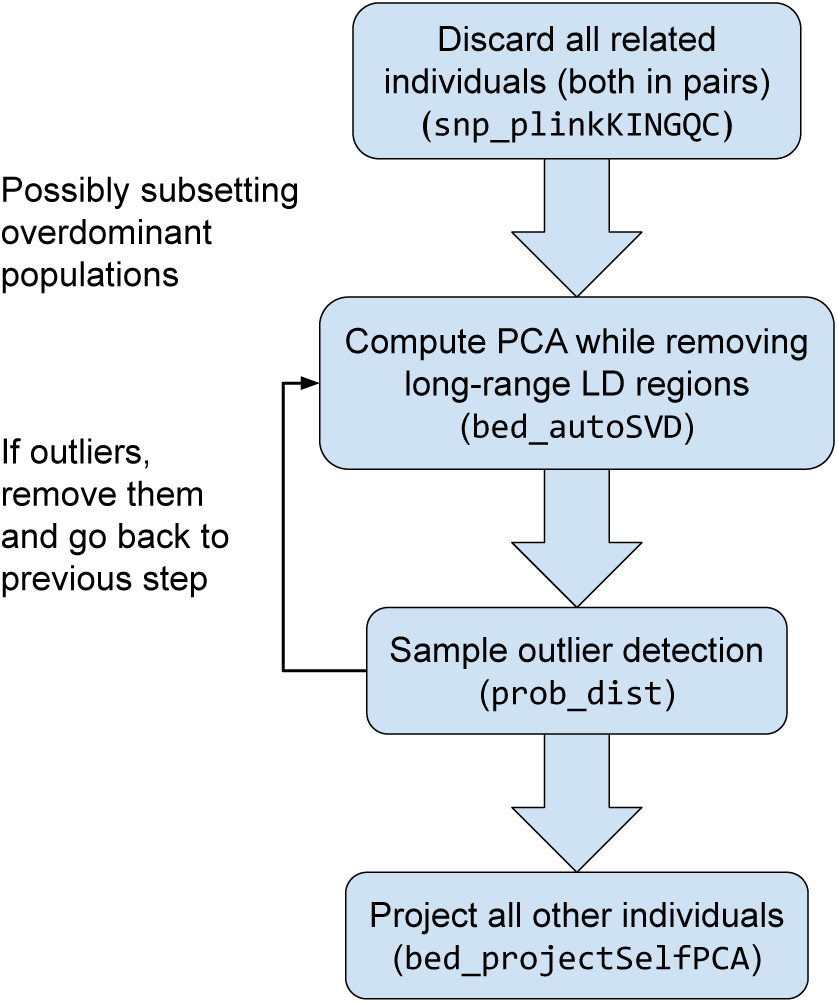
Proposed pipeline for computing Principal Components (PCs) using R packages bigsnpr and bigutilsr.

Once PCA is done, one should check the PC scores (scores of each individual for each PCA dimension) and PC loadings (weights for each variant for each PCA dimension). We differentiate PCs in 3 broad types: the ones capturing LD structure, the ones capturing population structure, and noise. We expect population structure to be well distributed along the genome so that we get loadings that are normally distributed around 0 (with small effect sizes). In contrast, long-range LD structure is basically capturing the variation inside one long-range LD region (so localised in the genome), so that we expect the loadings to be very large in that region only (one peak). Therefore, PCs capturing LD structure can be identified by looking for peaks in PC loadings (e.g. PC17-20 in figure S10). To identify which PCs capture population structure, and which ones are probably just noise, one should also look at PC scores (colored by ancestry if possible). As in many applications, we believe a compromise between signal and noise should be preferred. Therefore, we recommend using only PCs that show structure (e.g. PC1-16 in figure S9) and excluding PCs that do not seem to capture any population structure (e.g. PC17-20 in figure S9).

When analysing a dataset that is composed mainly of one population (e.g. British people in the UK Biobank), we found that it is useful to subset these individuals to reduce the imbalance between the different population sizes. Likewise, previous works have shown that uneven population sizes can distort PCs (Novembre and Stephens 2008; McVean 2009). Indeed, when subsetting British and Irish people in the UK Biobank data, we are able to capture a lot more PCs that show population structure with less than 50K individuals as compared to when using more than 400K individuals who are mostly composed of British and Irish people. Determining how much overdominant populations should be subsampled to maximise population structure captured by PCA is a direction of future work. The remaining individuals can then be projected onto the resulting PCA space using the OADP projection we recommend in this paper. This suggests that designs such as the 1000 Genomes project, which gathered around 100 people for each of 26 different populations, are highly relevant for capturing population structure (1000 Genomes Project Consortium *et al.* 2015).

In contrast, a common strategy in genetic analyses is to restrict the analysis to a homogeneous sample to reduce risk of confounding due to population stratification. For that purpose, we show that using the Mahalanobis distance on PC scores can efficiently achieve this goal, which we used in previous analyses (Privé *et al.* 2019). When the homogeneous sample is not predominant in the dataset, one solution is to compute the center and covariance of the robust Mahalanobis distance using only the population of interest, and then computing the distances for all individuals using these robust estimates.

The ubiquitous use of PCA in a wide variety of genomic analyses makes it difficult to establish universal guidelines for such analysis. Although we have tackled many problems related to computing PCA on genotype data in this paper, we do not answer other important problems, such as how to best control for population structure in genomic analyses. For example, when conducting a GWAS, should one restrict to a homogenous sample, or is it enough to just include PCs that capture population structure as covariates, or should one also use PCs as covariates in mixed linear models (Price *et al.* 2010; Loh *et al.* 2015b)? Similarly, in some analyses, it may be beneficial to include PCs that capture long-range or even inter-chromosomal LD. More work is needed to understand these fundamental problems, and to provide precise guidelines for conducting successful GWAS, heritability, and other genomic analyses where PCA is used. These are directions of future work.

Finally, although we have focused on PCA of genotype data in this paper, we believe most of the results presented here are not inherent to genotype data, and can be transferred to e.g. other omics data as well. For example, PCs can be used to account for confounding in other data as well (Pickrell *et al.* 2010). Then, outlier and homogeneous sample detection can be used on PCs of other types of data. Moreover, projection of scores will also be a problem for other omics data where the number of variables used is larger than the number of samples used for computing PCA. Finally, using “populations” with approximately the same size is relevant for other biological data as well. However, other pitfalls might apply when using other types of data; for example, methylation data can be confounded by factors such as age and sex, and it might be beneficial to remove some methylation probes that are associated with these confounding factors before computing PCA (Decamps *et al.* 2019).

## Software and code availability

All code used for this paper is available at https://github.com/privefl/paper4-bedpca/tree/master/code. Package bigsnpr can be installed from GitHub (see https://github.com/privefl/bigsnpr). A tutorial on the steps to perform PCA on 1000G data is available at https://privefl.github.io/bigsnpr/articles/bedpca.html.

## Acknowledgements

Authors thank Rounak Dey and Seunggeun Lee for helpful discussions about PCA projection. This research has been conducted using the UK Biobank Resource under Application Number 25589.

F.P., J.M. and B.V. are supported by the Danish National Research Foundation (Niels Bohr Professorship to J.M.), and also acknowledge the Lundbeck Foundation Initiative for Integrative Psychiatric Research, iPSYCH (R248-2017-2003).

## Supplementary Materials

### Optimised OADP transformation

We implement an optimised version of the Online Augmentation, Decomposition, and Procrustes (OADP) transformation when using *K*″ = *K*′ = *K* (Zhang *et al.* 2019). We assume that the *K*-partial Singular Value Decomposition (SVD) of the reference matrix *X* (of size *n* × *p*) has been computed as *U* Δ*V* ^*T*^. There are several steps to perform OADP transformation for each sample *y* (of size 1 × *p*) of target matrix *Y* (of size *m* × *p*):

1. Calculate *l* = *y*·*V* (of size 1×*K*), where *V* are the K PC loadings. And *g* = *y*·*h*^*T*^ (of size 1×1), where *h* = (*y*−*l*·*V* ^*T*^)*/*‖*y*−*l*·*V* ^*T*^ ‖_2_. Actually, 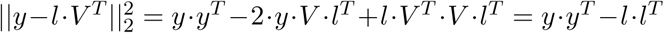 and *y* · (*y* − *l* · *V* ^*T*^)^*T*^ = *y* · *y*^*T*^ − *y* · *V* · *l*^*T*^ = *y* · *y*^*T*^ − *l* · *l*^*T*^. Then 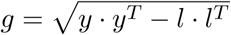.
2. Calculate *Q*^*T*^ *Q* where 

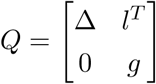

so that

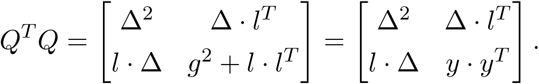 Note that we do not actually need to compute *g*, and that we can update only the last row and column of *Q*^*T*^ *Q* instead of computing it from an updated version of *Q*.
3. Get the eigen decomposition *Q*^*T*^ *Q* = *V* ′Δ′^2^*V* ′^*T*^ (truncated to *K* components out of the *K* + 1). Let us denote *V*_2_ = *V* ′Δ′.
4. Calculate

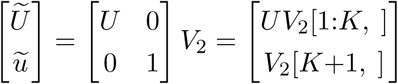
5. Find the Procrustes transformation from 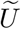 to *U* Δ. As 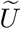 and *U* Δ have both their columns centered already (since *U* does), the Procrustes transformation 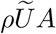, where *ρ* is a scaling coefficient and *A* is an orthonomal projection matrix that minimise the Frobenius norm of 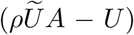, is given by *A* = *V* ″ *U* ″^*T*^and 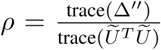 where *U* ″Δ″*V* ″^*T*^ is the full SVD of 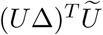 (Wang *et al.* 2015). As *U*^*T*^ *U* = *I*, we note that 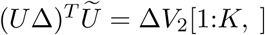 and that 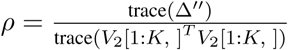, therefore we do not need to explicitly compute 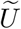 and do not need *U*.
6. Apply the previous transformation to 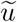 to get the projection of *y* in the reference PCA space (i.e. 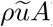).

### Sample outlier detection

**Figure S1:**
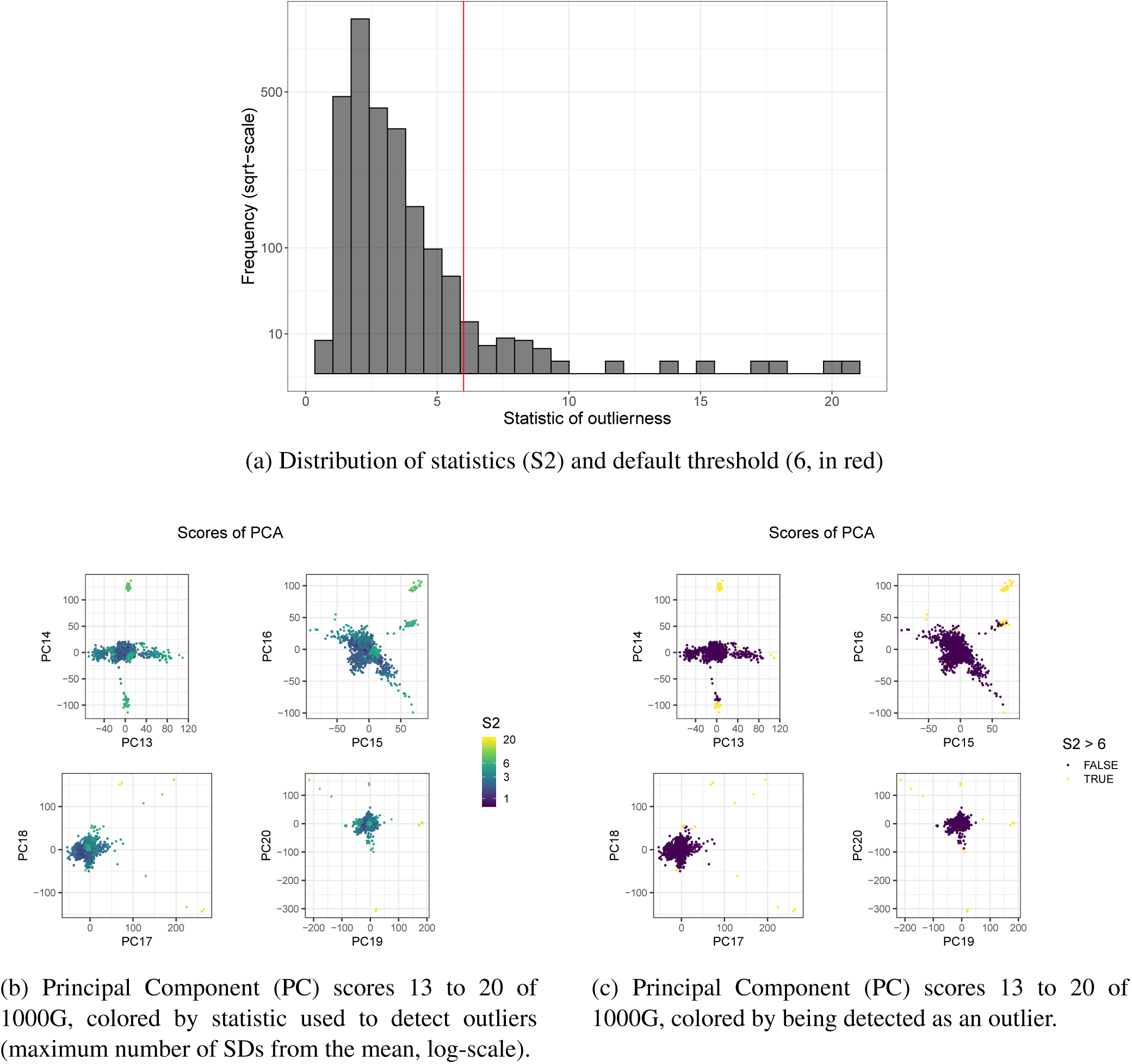
Outlier detection in the 1000 Genomes (1000G) project, using the rule “6 SDs from the mean”, i.e. where S2 is the maximum (for all PCs) number of SDs from the mean (Section 2.4).

**Table S1:**
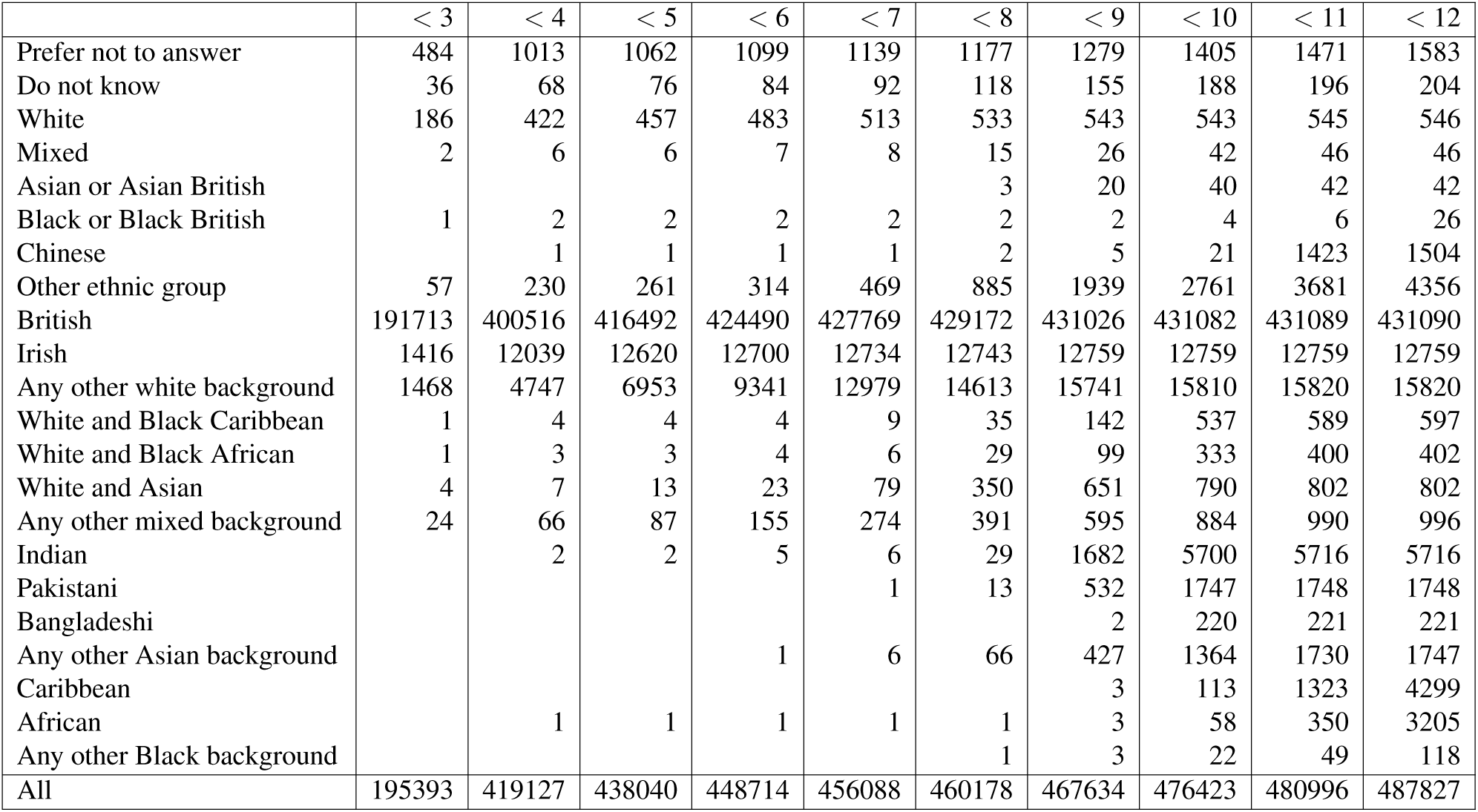
Number of UKBB individuals with (log squared) Mahalanobis distance lower than some threshold (top), and grouped by self-reported ancestry (left). Note that “< 12” includes all individuals.

**Figure S2:**
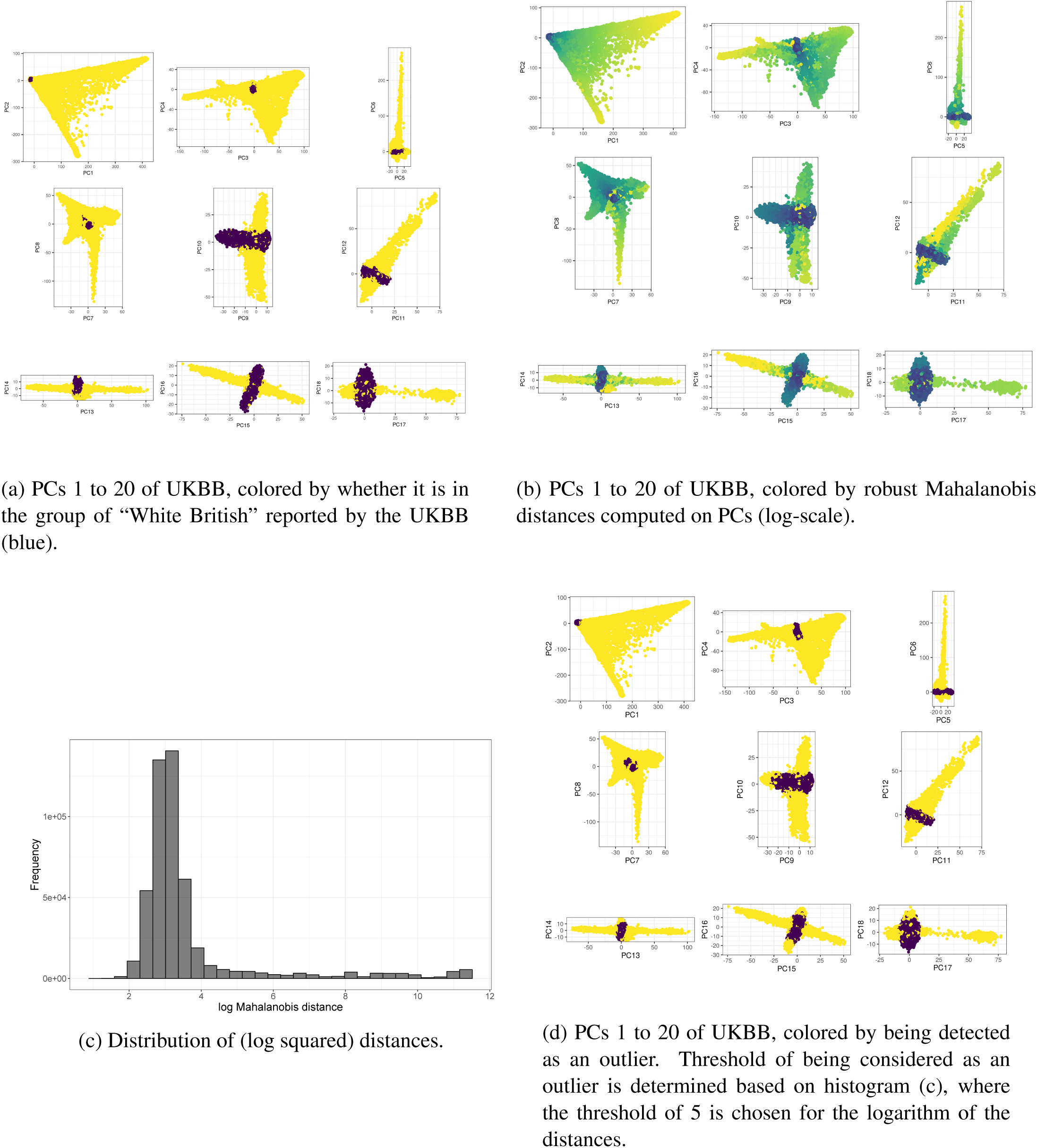
Homogeneous sample detection in the UK Biobank (UKBB), using robust Mahalanobis distances computed on the first 20 Principal Component scores (PCs) of UKBB.).

### Projection onto reference PCA space

**Figure S3:**
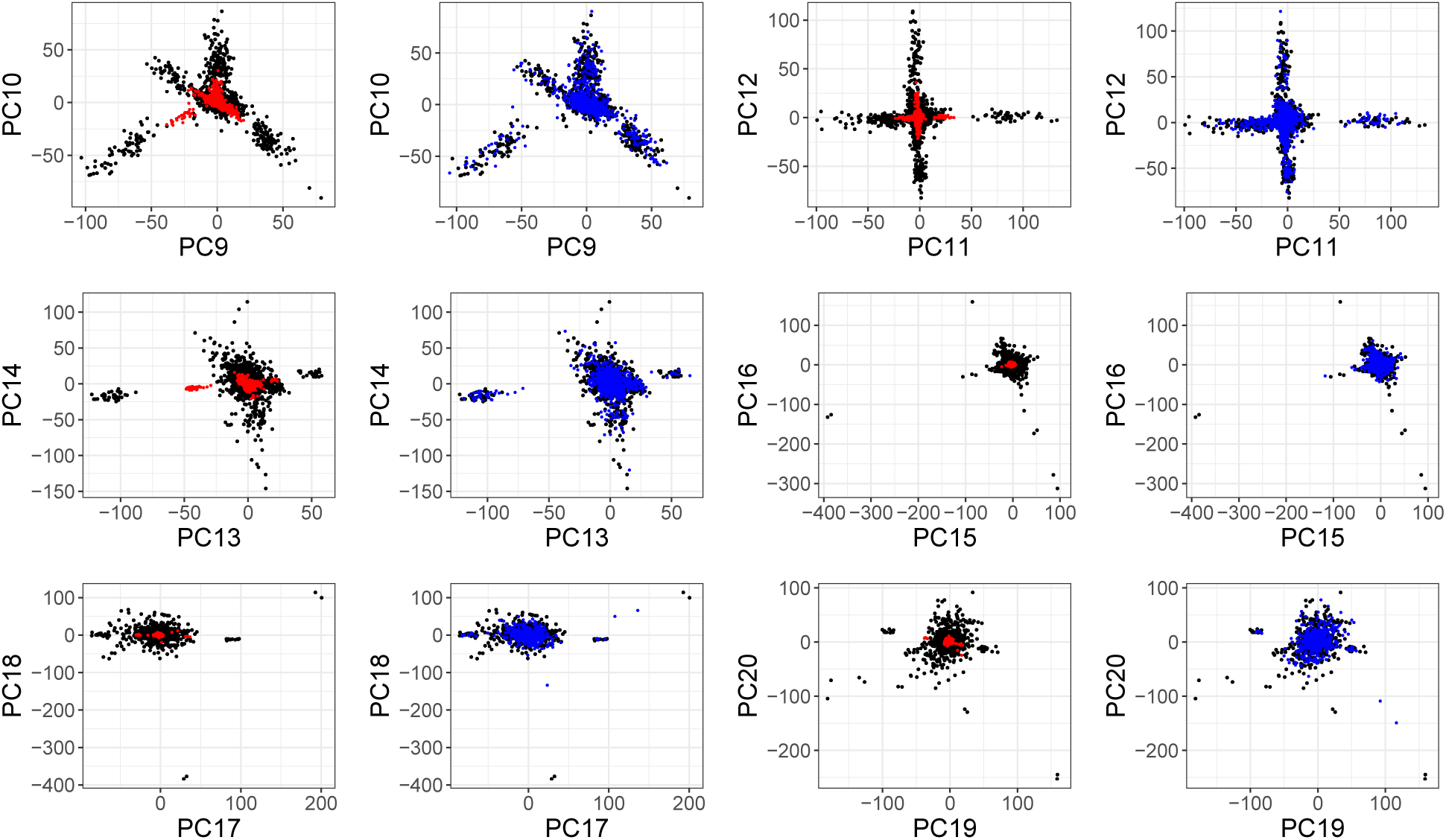
Principal Component (PC) scores 9 to 20 of the 1000 Genomes project. Black points are the 60% individuals used for computing PCA. Red points are the 40% remaining individuals, projected by simply multiplying their genotypes by the corresponding PC loadings. Blue points are the 40% remaining individuals, projected using the Online Augmentation, Decomposition, and Procrustes (OADP) transformation. Estimated shrinkage coefficients (comparing red and blue points) for these PCs are 2.79, 3.14 (PC10), 3.64, 3.18, 2.47, 3.88, 5.31, 5.84, 3.45, 6.55, 3.68 and 6.70 (PC20).

**Figure S4:**
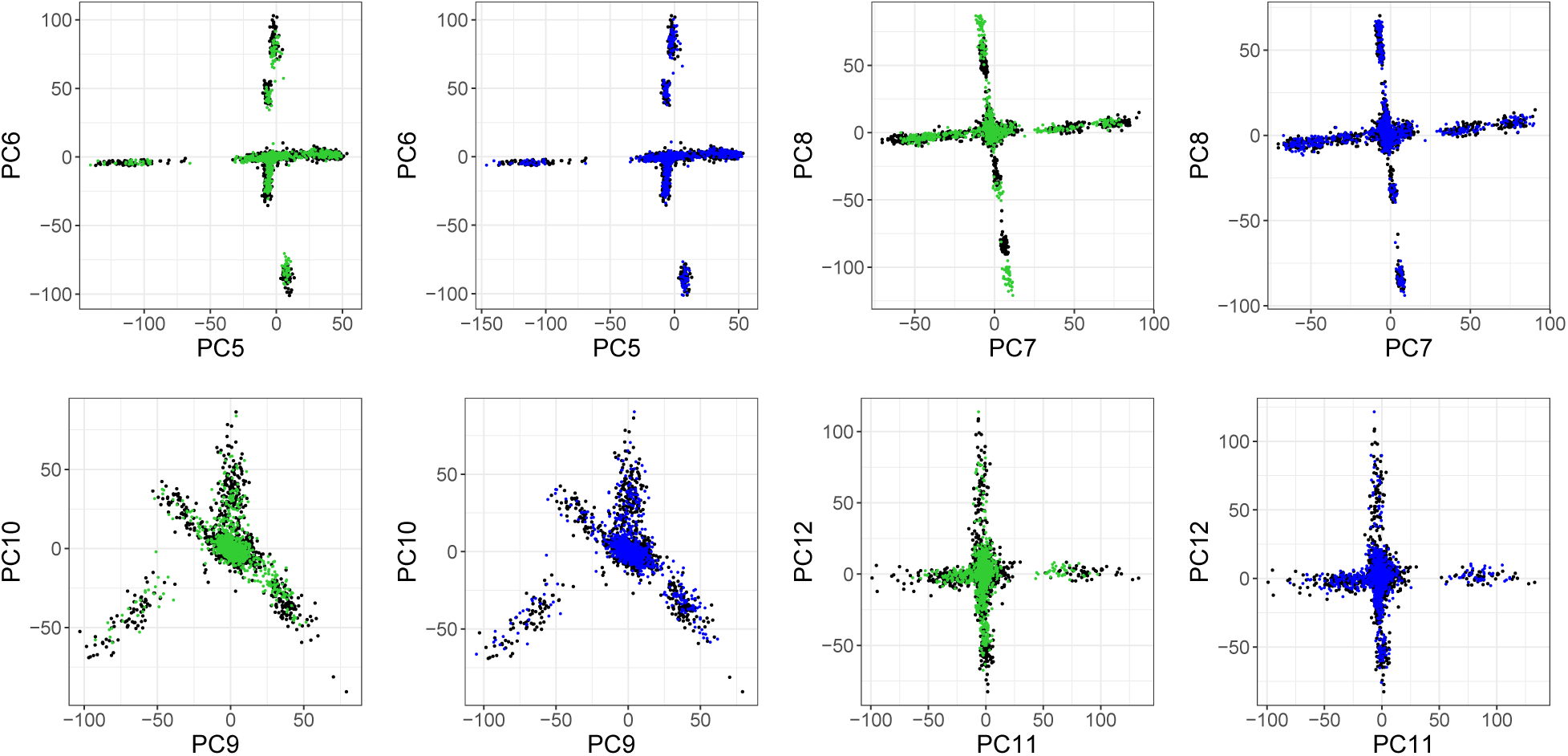
Principal Component (PC) scores 5 to 12 of the 1000 Genomes project. Black points are the 60% individuals used for computing PCA. Green points are the 40% remaining individuals, projected by multiplying their genotypes by the corresponding PC loadings, further corrected using theoritical asymptotic shrinkage factors (values for the first 12 PCs: 1.01 (PC1), 1.02, 1.07, 1.10, 1.43 (PC5), 1.54, 1.74, 1.79, 2.47, 2.77 (PC10), 2.84 and 3.15). Blue points are the 40% remaining individuals, projected using the Online Augmentation, Decomposition, and Procrustes (OADP) transformation.

**Figure S5:**
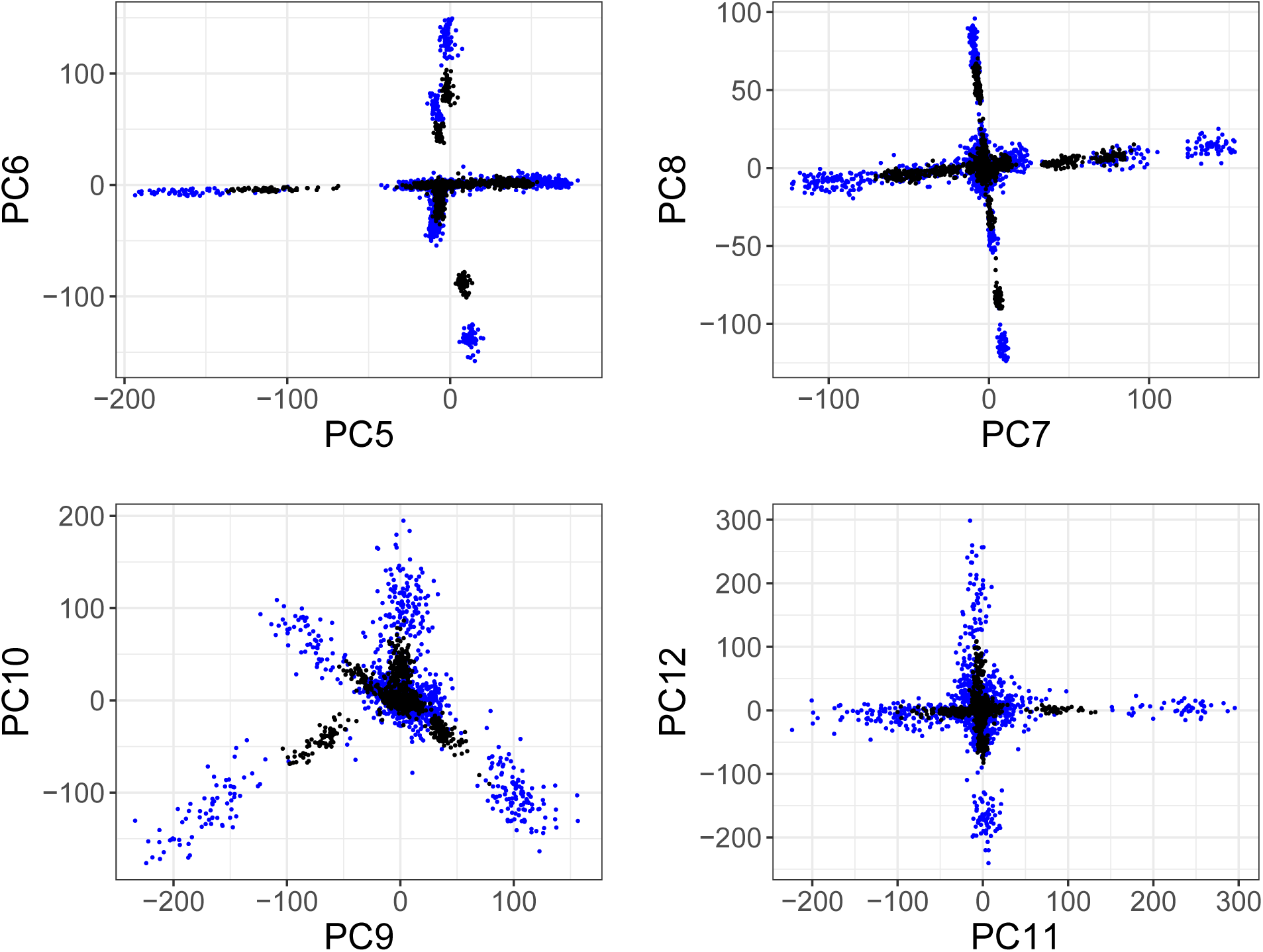
Principal Component (PC) scores 5 to 12 of the 1000 Genomes project. Black points are the 60% individuals used for computing PCA. Blue points are the same 60% individuals, projected using the Online Augmentation, Decomposition, and Procrustes (OADP) transformation.

**Figure S6:**
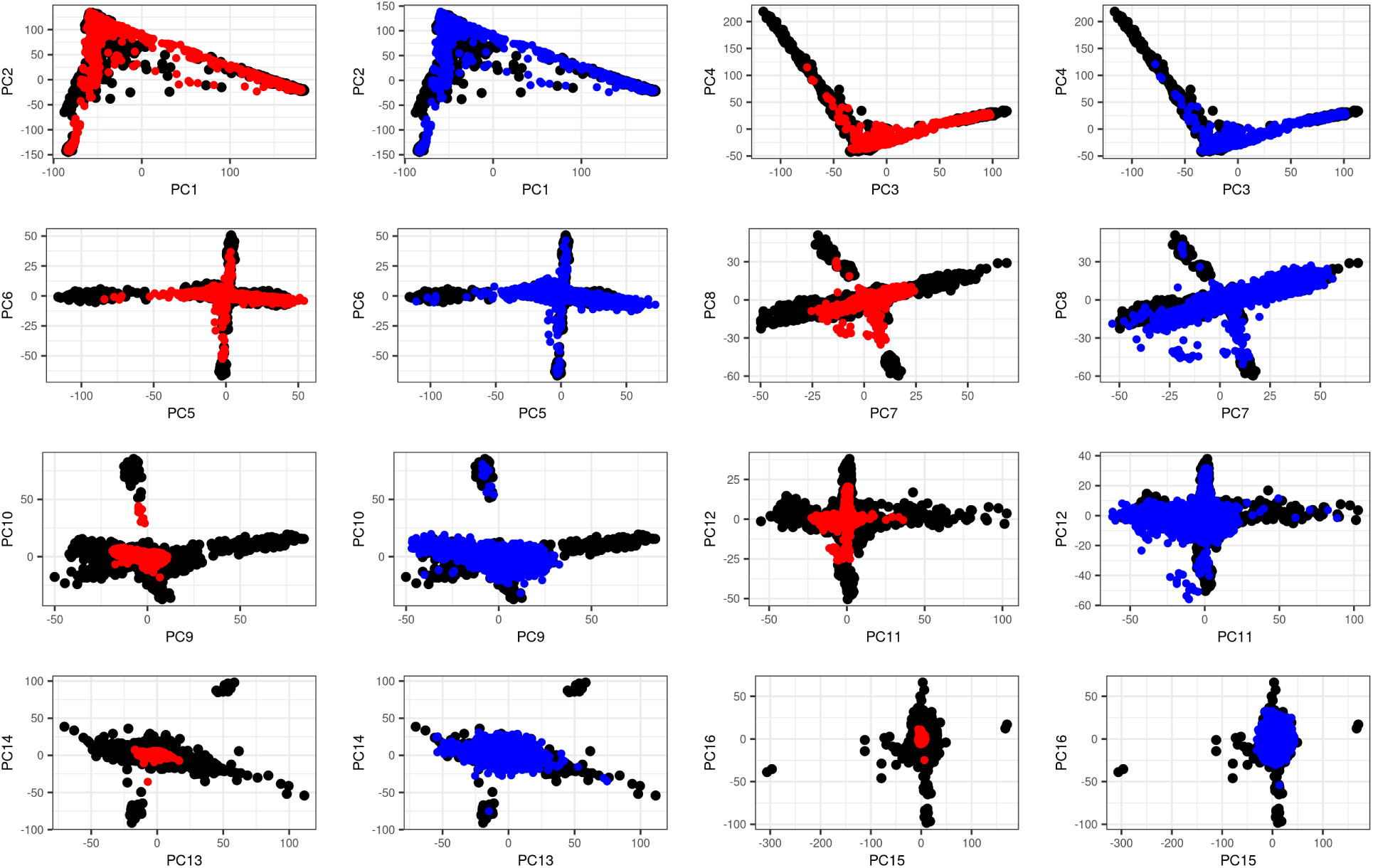
Principal Component (PC) scores 1 to 16 of the 1000 Genomes project and projected individuals from the UK Biobank. Black points are PC scores of 1000G individuals used for computing PCA. Red points are the individuals from UKBB, projected by simply multiplying their genotypes by the corresponding PC loadings. Blue points are the 488,371 individuals from the UK Biobank, projected using the Online Augmentation, Decomposition, and Procrustes (OADP) transformation. Estimated shrinkage coefficients (comparing red and blue points) for the first 20 PCs are 1.01 (PC1), 1.02, 1.06, 1.08, 1.36 (PC5), 1.82, 2.33, 2.36, 2.78, 2.84 (PC10), 2.99, 3.51, 4.38, 4.67, 4.99, 5.31, 5.74, 6.55, 6.71 and 6.75 (PC20). Note that only 20,000 random projected individuals are represented in this plot.

**Figure S7:**
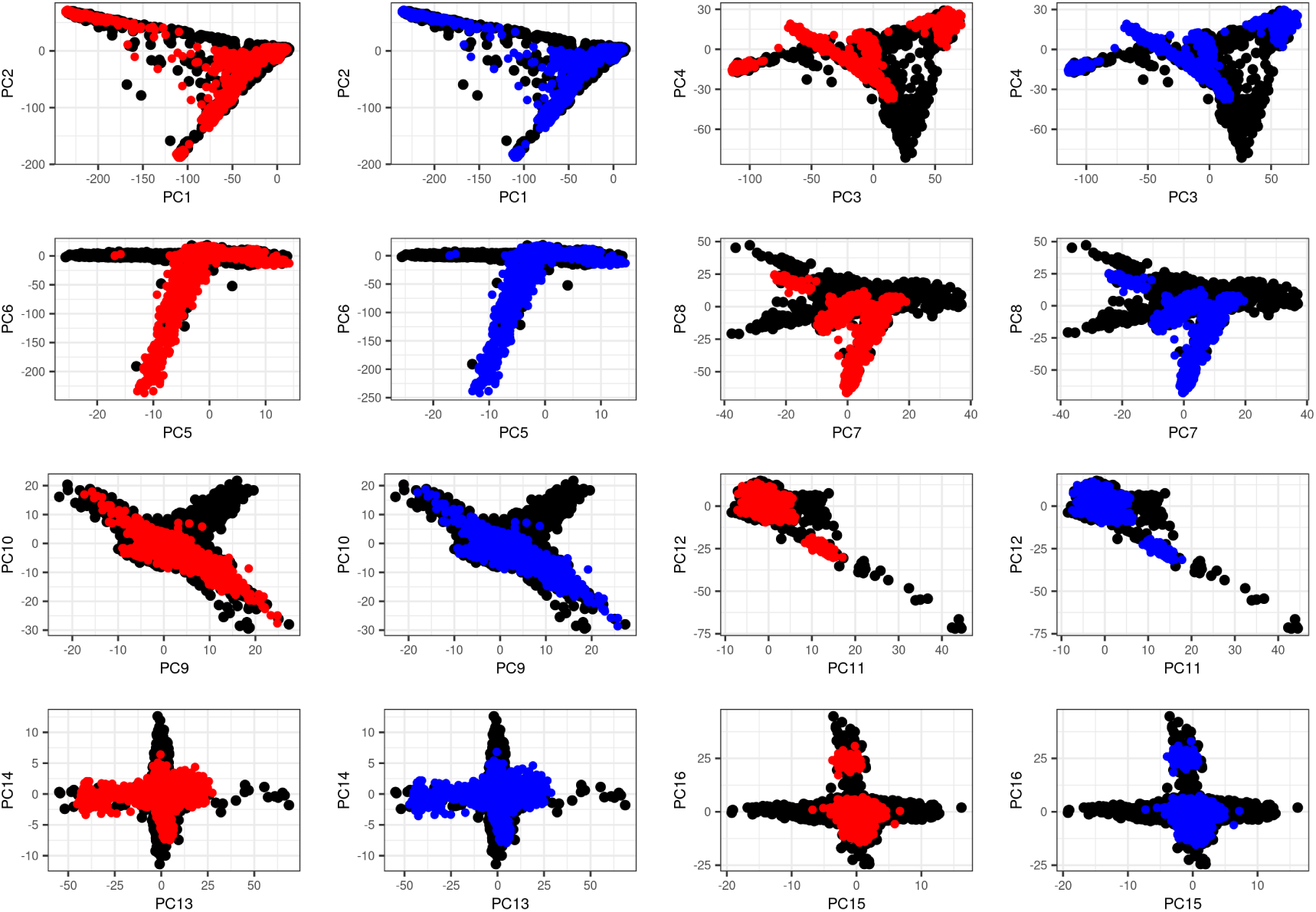
Principal Component (PC) scores 1 to 16 from the UK Biobank and projected individuals of the 1000 Genomes (1000G) project. Black points are the UK Biobank individuals used for computing PCA. Red points are the individuals from 1000G, projected by simply multiplying their genotypes by the corresponding PC loadings. Blue points are the individuals from 1000G, projected using the Online Augmentation, Decomposition, and Procrustes (OADP) transformation. Estimated shrinkage coefficients (comparing red and blue points) for the first 20 PCs are 1.00 (PC1), 1.00, 1.00, 1.01, 1.01 (PC5), 1.02, 1.03, 1.03, 1.04, 1.04 (PC10), 1.04, 1.05, 1.05, 1.06, 1.07, 1.07, 1.08, 1.08, 1.08 and 1.08 (PC20).

### PCA of the UK Biobank

**Figure S8:**
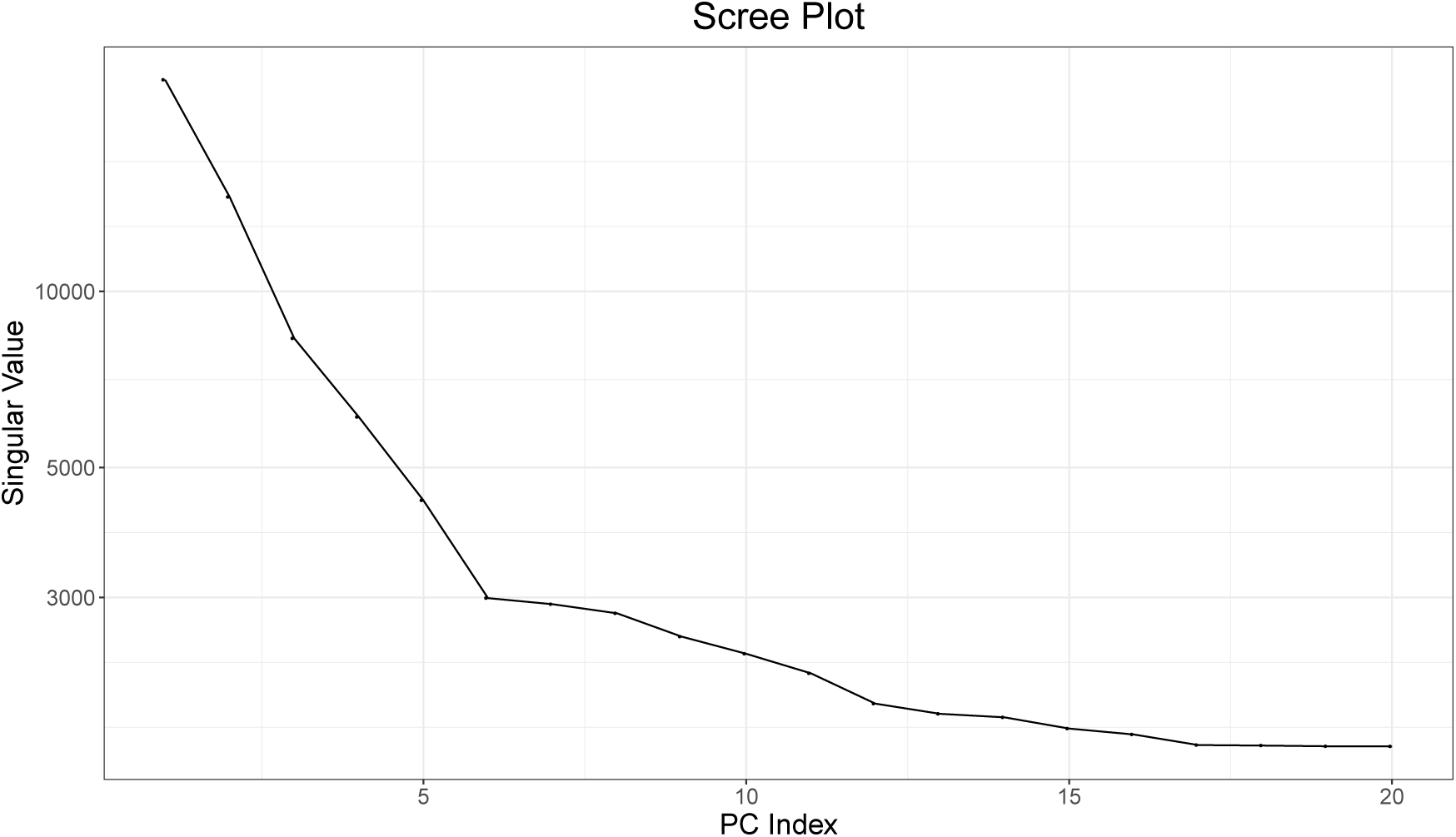
Scree plot: plot of singular values computed on the UK Biobank using `bed_autoSVD`.

**Figure S9:**
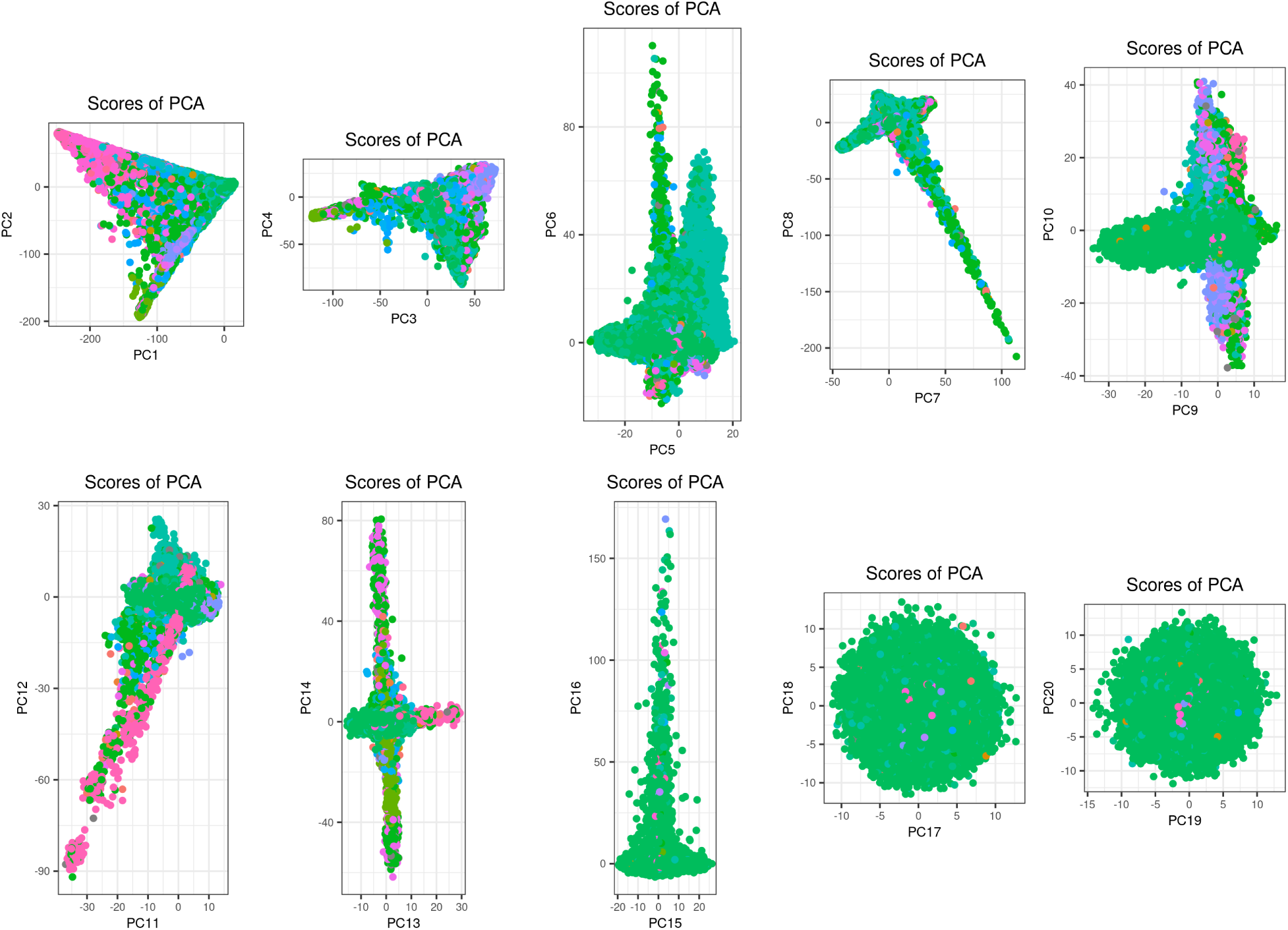
Principal Component (PC) scores 1 to 20 computed on the UK Biobank using `bed_autoSVD`. Different colors represent different self-reported ancestries.

**Figure S10:**
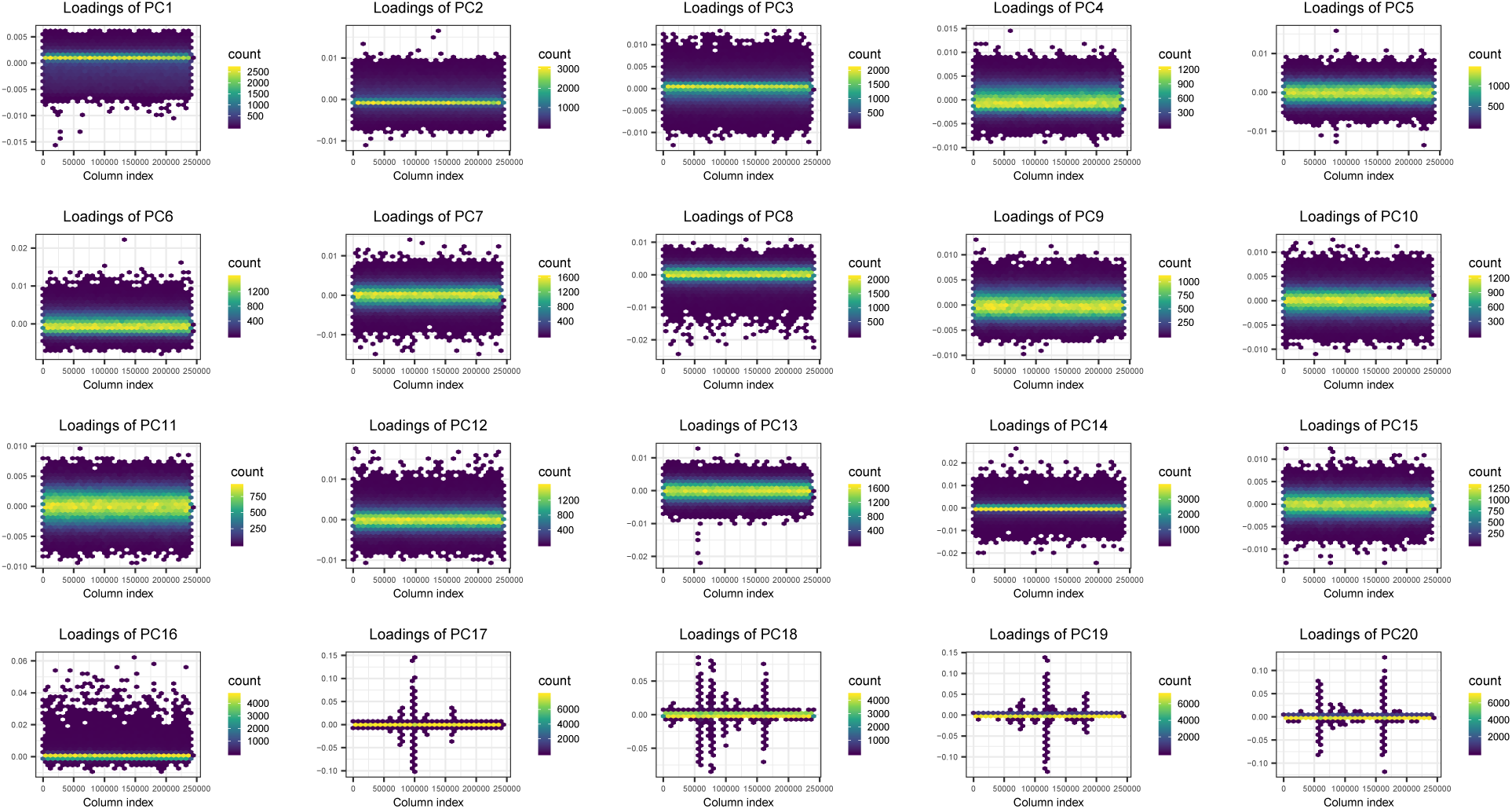
Principal Component (PC) loadings 1 to 20 computed on the UK Biobank using `bed_autoSVD`. X-axis represent positions of variants in the data, which is ordered by chromosome and physical position, and y-axis the value of loadings. Points are hex-binned to make plotting of such large number of points manageable.

**Figure S11:**
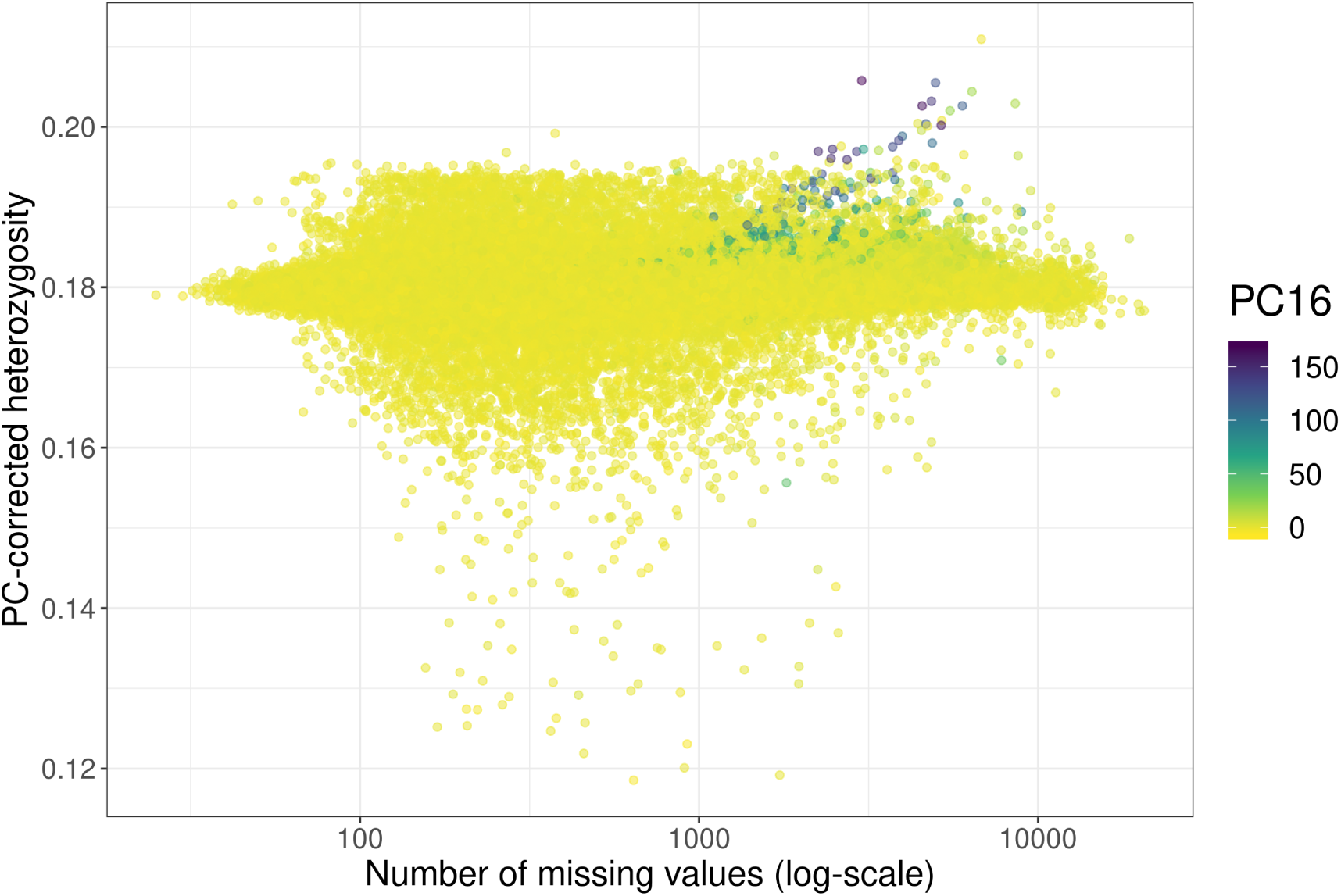
PC-corrected heterozygosity and number of missing values for individuals of the UK Biobank, colored by their value for PC16.

**Table S2:** Correlation between first 20 PC scores of the UK Biobank, both computed using the genotyping chip and mean imputation, but wtih either no quality control on individuals based on high heterozygosity (top), or after removing some individuals based on high heterozygosity (left). PC16 computed without quality control on high heterozygosity completely disappears when performing this quality control.

|  | 1 | 2 | 3 | 4 | 5 | 6 | 7 | 8 | 9 | 10 | 11 | 12 | 13 | 14 | 15 | 16 | 17 | 18 | 19 | 20 |
| --- | --- | --- | --- | --- | --- | --- | --- | --- | --- | --- | --- | --- | --- | --- | --- | --- | --- | --- | --- | --- |
| 1 | 100.0 | -0.2 | 0.1 | 0.0 | 0.0 | 0.0 | 0.0 | 0.0 | -0.0 | -0.0 | 0.0 | 0.0 | -0.0 | -0.0 | -0.0 | -0.3 | -0.0 | 0.0 | 0.0 | 0.0 |
| 2 | 0.1 | 100.0 | 0.2 | 0.0 | 0.0 | 0.0 | 0.0 | 0.0 | 0.0 | -0.1 | 0.0 | -0.0 | 0.0 | -0.0 | -0.0 | 0.0 | -0.0 | -0.0 | 0.0 | 0.0 |
| 3 | -0.0 | -0.1 | 100.0 | -0.0 | 0.0 | -0.0 | -0.0 | -0.0 | -0.0 | 0.1 | -0.0 | 0.0 | 0.0 | -0.0 | -0.0 | -0.3 | 0.0 | 0.0 | -0.0 | -0.0 |
| 4 | -0.0 | -0.0 | 0.0 | 100.0 | -0.0 | 0.0 | 0.0 | 0.0 | -0.0 | -0.0 | 0.0 | 0.0 | -0.0 | -0.0 | -0.0 | 0.1 | 0.0 | 0.0 | 0.0 | 0.0 |
| 5 | 0.0 | -0.0 | -0.0 | 0.0 | 100.0 | 0.0 | -0.0 | 0.0 | 0.0 | -0.0 | -0.0 | 0.0 | 0.0 | 0.0 | -0.0 | 0.1 | -0.0 | -0.0 | -0.0 | 0.0 |
| 6 | -0.0 | -0.0 | 0.0 | -0.0 | -0.0 | 100.0 | -0.3 | 0.4 | -0.0 | 0.0 | -0.0 | 0.0 | -0.0 | -0.0 | 0.0 | -0.1 | 0.0 | -0.0 | -0.0 | 0.0 |
| 7 | -0.0 | -0.0 | 0.0 | -0.0 | 0.0 | 0.3 | 100.0 | -0.2 | -0.0 | 0.2 | -0.1 | -0.1 | 0.0 | 0.0 | 0.0 | 0.3 | -0.0 | -0.0 | 0.0 | -0.0 |
| 8 | -0.0 | -0.0 | 0.0 | -0.0 | -0.0 | -0.4 | 0.2 | 100.0 | 0.0 | 0.2 | -0.1 | -0.0 | 0.0 | 0.0 | 0.0 | 0.2 | 0.0 | 0.0 | -0.0 | 0.0 |
| 9 | 0.0 | -0.0 | 0.0 | 0.0 | -0.0 | 0.0 | 0.0 | -0.0 | 100.0 | 0.2 | -0.0 | 0.0 | -0.0 | -0.0 | 0.0 | -0.5 | -0.0 | 0.0 | -0.0 | 0.0 |
| 10 | 0.0 | 0.0 | -0.0 | 0.0 | 0.0 | -0.0 | -0.1 | -0.1 | -0.2 | 100.0 | 0.4 | -0.1 | 0.1 | 0.0 | 0.0 | 0.8 | -0.0 | -0.0 | 0.0 | 0.0 |
| 11 | -0.0 | -0.0 | 0.0 | -0.0 | 0.0 | 0.0 | 0.0 | 0.0 | 0.0 | -0.3 | 100.0 | 0.2 | -0.1 | -0.0 | -0.0 | -0.2 | 0.0 | 0.0 | -0.0 | 0.0 |
| 12 | -0.0 | 0.0 | -0.0 | -0.0 | -0.0 | -0.0 | 0.0 | 0.0 | -0.0 | 0.1 | -0.1 | 100.0 | -0.3 | 0.1 | 0.0 | 1.7 | -0.0 | -0.1 | -0.0 | -0.0 |
| 13 | 0.0 | -0.0 | -0.0 | 0.0 | -0.0 | 0.0 | -0.0 | -0.0 | 0.0 | -0.0 | 0.0 | 0.3 | 100.0 | -0.4 | -0.1 | -2.8 | 0.0 | 0.0 | 0.1 | 0.0 |
| 14 | 0.0 | 0.0 | 0.0 | 0.0 | -0.0 | 0.0 | -0.0 | -0.0 | -0.0 | -0.0 | 0.0 | -0.1 | 0.4 | 100.0 | -0.0 | -1.8 | 0.0 | 0.0 | 0.0 | -0.0 |
| 15 | 0.0 | 0.0 | -0.0 | 0.0 | 0.0 | -0.0 | 0.0 | -0.0 | -0.0 | -0.0 | 0.0 | -0.0 | 0.0 | -0.0 | 100.0 | -2.1 | 0.1 | -0.0 | 0.0 | 0.0 |
| 16 | 0.0 | 0.0 | -0.0 | -0.0 | 0.0 | -0.0 | 0.0 | -0.0 | 0.0 | 0.0 | -0.0 | 0.0 | 0.0 | 0.0 | -0.0 | 1.3 | 100.0 | 0.6 | -0.1 | -0.3 |
| 17 | -0.0 | -0.0 | -0.0 | 0.0 | -0.0 | -0.0 | -0.0 | 0.0 | -0.0 | -0.0 | 0.0 | 0.0 | -0.0 | -0.0 | -0.1 | -5.0 | 0.7 | -99.9 | -0.8 | -0.2 |
| 18 | 0.0 | 0.0 | -0.0 | 0.0 | -0.0 | -0.0 | 0.0 | 0.0 | -0.0 | 0.0 | -0.0 | 0.0 | -0.0 | -0.0 | -0.0 | -2.9 | 0.0 | 0.9 | -99.5 | 8.7 |
| 19 | 0.0 | 0.0 | -0.0 | 0.0 | 0.0 | 0.0 | -0.0 | 0.0 | 0.0 | 0.0 | 0.0 | -0.0 | 0.0 | -0.0 | 0.0 | -0.3 | -0.2 | 0.3 | -8.6 | -99.6 |
| 20 | 0.0 | 0.0 | -0.0 | 0.0 | -0.0 | -0.0 | -0.0 | 0.0 | 0.0 | 0.0 | 0.0 | -0.0 | 0.0 | 0.0 | 0.1 | 7.3 | -0.8 | -0.3 | -4.4 | -0.3 |

**Table S3:**
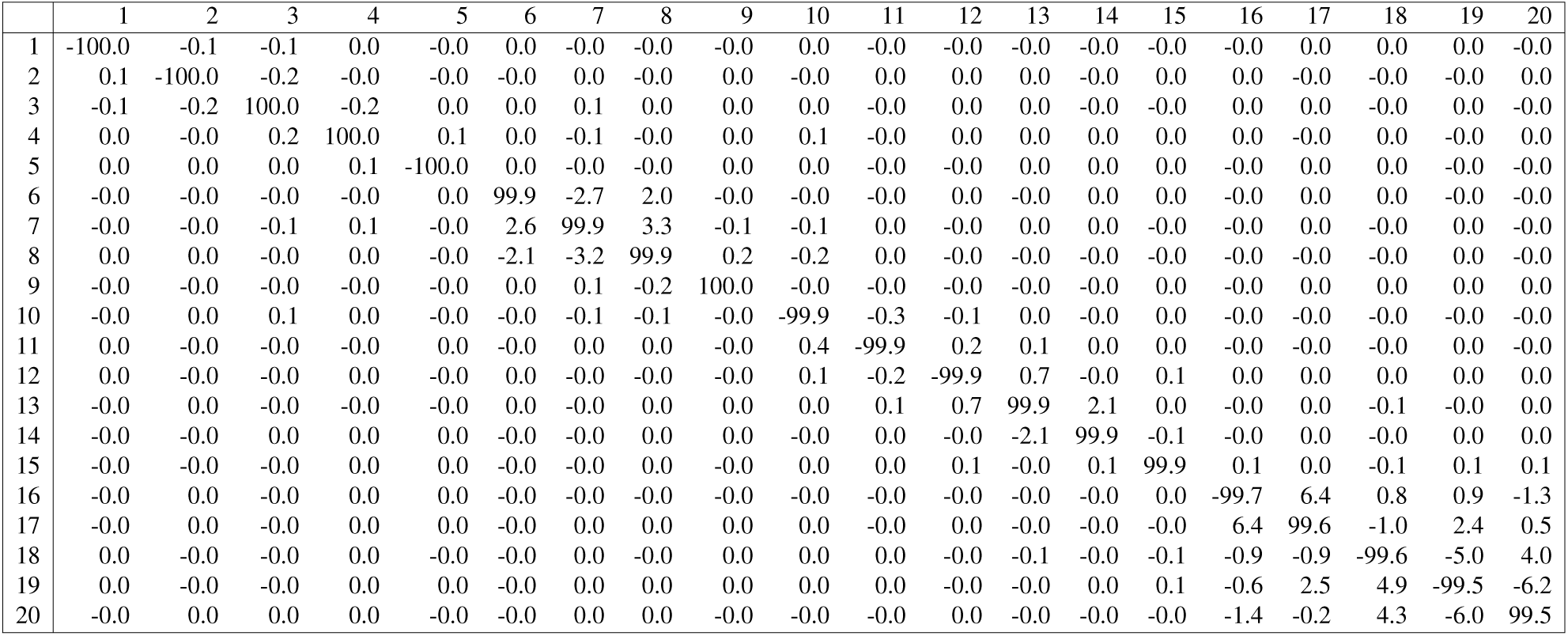
Correlation between first 20 PC scores of the UK Biobank, either computed using the genotyping chip and mean imputation (top), or computed from the dosages (based on imputation from an external reference panel) of the same variants (left). PCs are globally the same.

**Figure S12:**
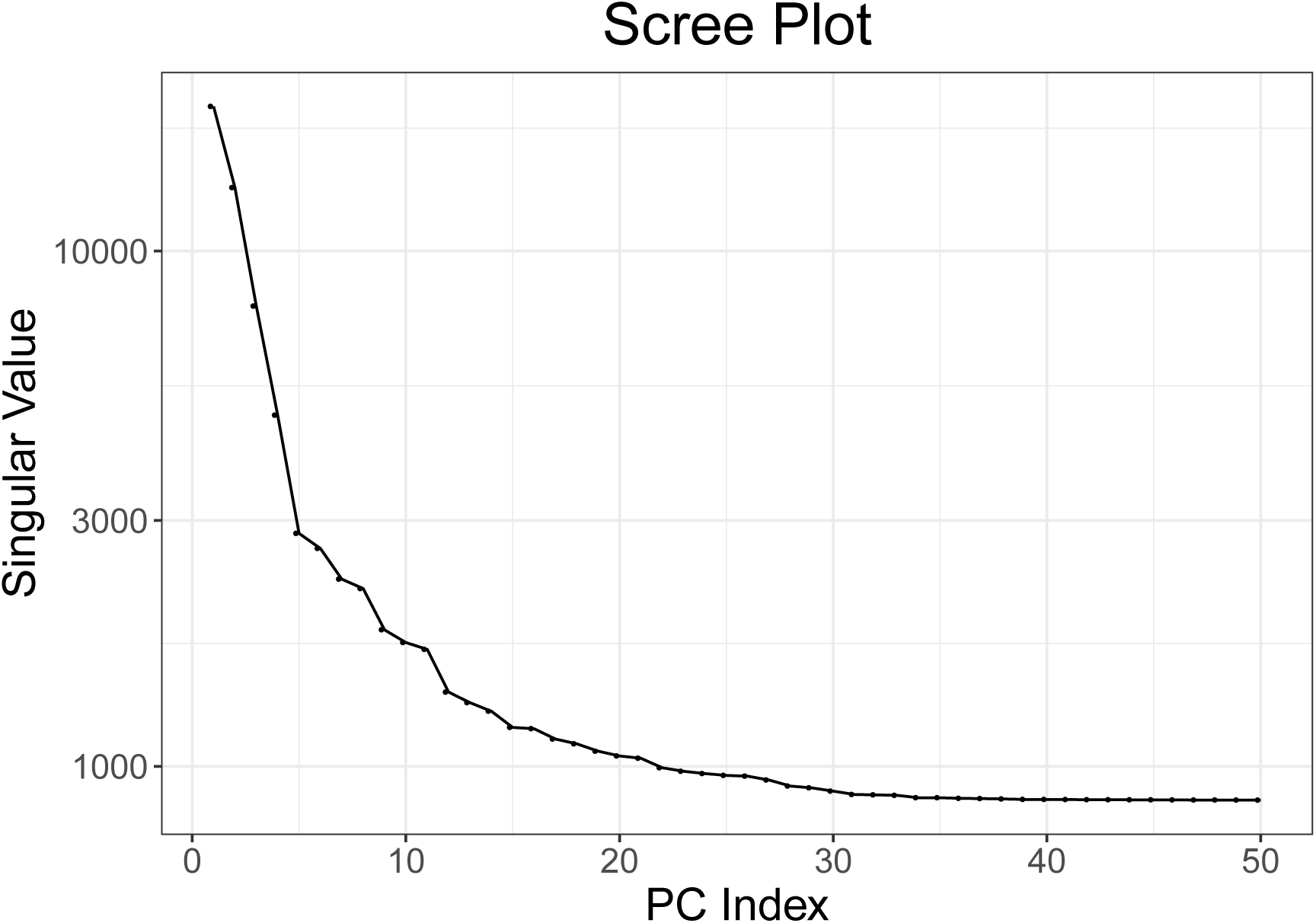
Scree plot: plot of singular values computed on the UK Biobank using 48,942 individuals of diverse ancestries. These individuals are the ones resulting from removing all related individuals and randomly subsampling the British and Irish individuals.

**Figure S13:**
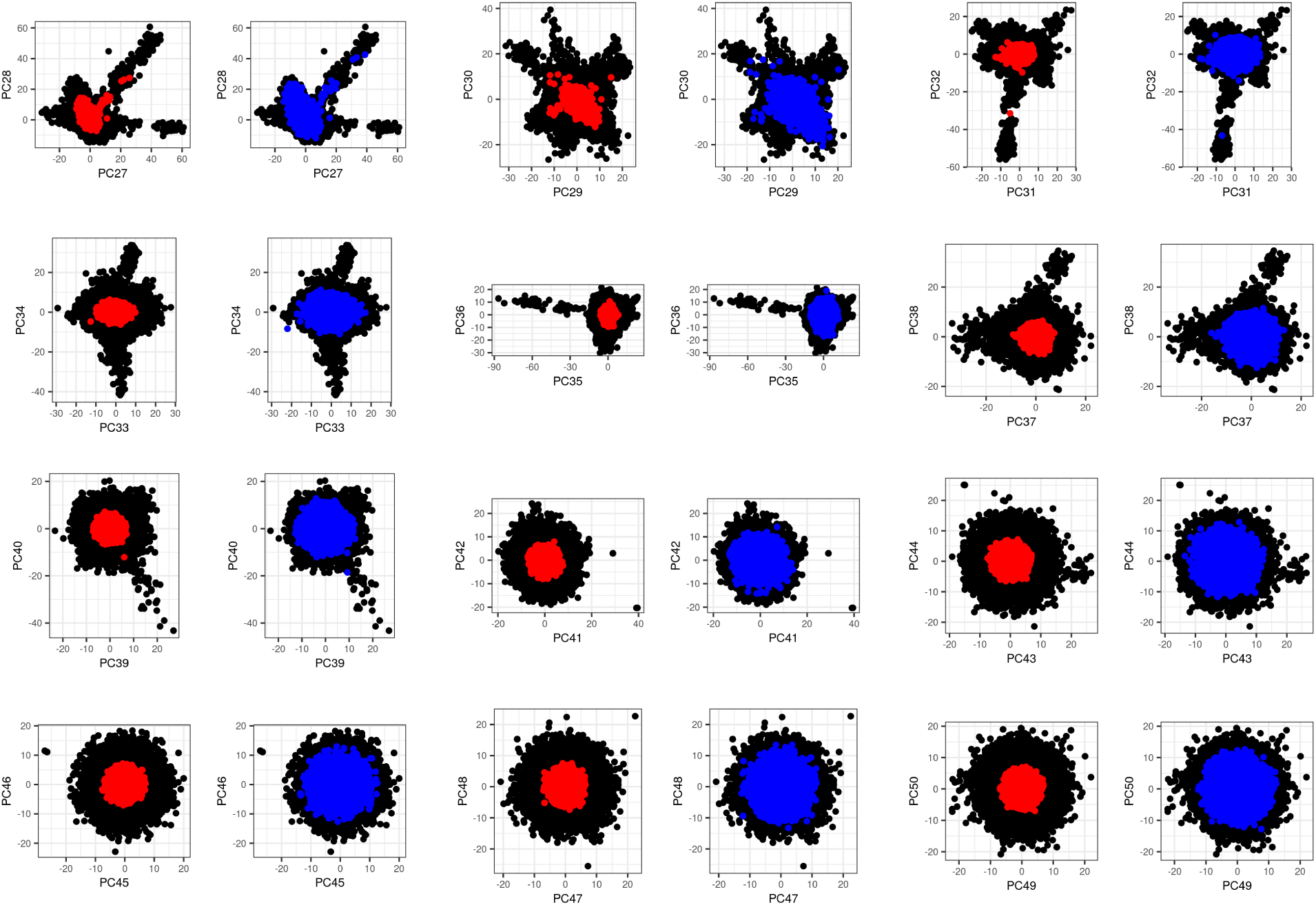
Principal Component (PC) scores 27 to 50 of the UK Biobank. Black points are the 48,942 individuals of diverse ancestries used for computing PCA. These individuals are the ones resulting from removing all related individuals and randomly subsampling the British and Irish individuals. Red points are the remaining UKBB individuals, projected by simply multiplying their genotypes by the corresponding PC loadings. Blue points are the remaining UKBB individuals, projected using the Online Augmentation, Decomposition, and Procrustes (OADP) transformation. Note that only 20,000 random projected individuals are represented in this plot.

